# Neural circuits between nodose ganglion and pulmonary neuroendocrine cells regulate lung inflammatory responses

**DOI:** 10.1101/2025.01.01.630820

**Authors:** Jie Chen, Shitao Xie, Zhekai Lin, Caiqi Zhao, Rujia Tao, Yingying Ma, Xiaoyan Chen, Renlan Wu, Pengfei Sui, Sheng Wang, Hongbin Ji, Hai Song, Xiaoming Zhang, Yangang Sun, Yuanlin Song, Xiao Su

**Affiliations:** Shanghai Institute of Immunity and infection, Chinese Academy of Sciences, University of Chinese Academy of Sciences, Shanghai 200031, China; Shanghai Key Laboratory of Lung Inflammation and Injury, Department of Pulmonary Medicine, Zhongshan Hospital, Fudan University, Shanghai 200032, China; Shanghai Institute of Biochemistry and Cell Biology, Chinese Academy of Sciences, Shanghai 200031, China; Department of Neurobiology, Hebei Medical University, Shijiazhuang, Hebei, China; The MOE Key Laboratory of Biosystems Homeostasis and Protection, Zhejiang Provincial Key Laboratory for Cancer Molecular Cell Biology and Innovation Center for Cell Signaling Network, Life Sciences Institute, Zhejiang University, Hangzhou, Zhejiang, 310058 China; Institute of Neuroscience, State Key Laboratory of Neuroscience, Center for Excellence in Brain Science & Intelligence Technology, Chinese Academy of Sciences, Shanghai, China; Key Laboratory of Immune Response and Immunotherapy, Shanghai Institute of Immunity and infection, Chinese Academy of Sciences, Shanghai 200031, China

**Keywords:** Vagus nerve, Nodose ganglion, LPS, Sensory neurons, Pathogen recognition, Lung inflammation, Pulmonary neuroendocrine cells (PNECs), Transient receptor potential ankyin1 (TRPA1), Alpha-Calcitonin gene related peptide (αCGRP)

## Abstract

The lungs are organs exposed to the external environment, and the air we inhale contains various pathogens, such as endotoxins. The vagus nerve, which innervates the lungs, may play a role in detecting pathogens that invade the lungs. Through transcriptome analysis, tissue clearance imaging, electrical excitability recording, and gene- and cell-specific knockout experiments, we found that vagus nerve endings innervate pulmonary neuroendocrine cells (PNECs). These nerve endings sense bacterial endotoxins via pain receptors (TRPA1) rather than toll-like receptors (TLR4), eliciting electrical excitation and enhancing the production of neuropeptides (αCGRP) in the nodose ganglia. In turn, αCGRP released by sensory neurons from the nodose ganglia promotes both neuropeptide production and the proliferation of PNECs, thereby amplifying endotoxin-induced lung inflammatory responses. This reveals that the neural circuits between the nodose ganglion and PNECs play a critical role in regulating lung inflammatory responses.

## Introduction

The jugular-nodose ganglia contain the sensory peripheral neurons of the vagus nerve, connecting visceral organs (i.e., lungs) to the medulla oblongata ^1–4^, and transmitting peripheral pathogenic information (i.e., lung lipopolysaccharide (LPS) assault) to the brain ^5,6^. In brain regions, caudal nucleus of the solitary tract (cNST) ^7^ and the paraventricular hypothalamic nucleus (PVN) ^8^ are activated to modulate the inflammatory response. The above-mentioned vagal modulation mechanism is named: parasympathetic inflammatory reflex ^6,9^. With regard to the efferent of parasympathetic inflammatory reflex in the lungs, we have reported that vagal-α7nAChR (nicotinic acetylcholine receptor) signaling regulates lung injury, viral replication, and stem cell repair ^10–13^. The afferents of this reflex have not been well studied.

As we know, the vagal sensory neurons form the jugular (*Prdm12*^+^) and nodose (*Phox2b*^+^) ganglia residing in the same nerve sheath ^1^. The nodose vagal ganglia innervate the distal airways of lung, while the jugular vagal ganglia innervate the proximal airway ^14^. Although TLR4 has been reported to detect LPS exposed in lung TRPV1^+^ sensory neurons ^8^, it remains unclear whether lung-innervated nodose sensory neurons express TLR4, which contributes to LPS perception in the lungs. In addition, LPS can activate suprarenal and celiac ganglia neurons and upregulate NPY expression by which affects the proliferation and activation of splenic lymphocytes ^15^. Similarly, we need to know if vagal sensory neurons can respond to lung LPS attack and increase the expression of their neuropeptides (i.e., CGRP) and regulate lung inflammation.

The lungs are an organ that is open to the outside environment. The air inhaled contains various pathogens (e.g., endotoxins). Neuro-epithelial crosstalk is crucial for sensing and regulating pulmonary inflammation. In the respiratory tract, sensory neurons serve as both first-order and second-order sensory neurons. They directly sense LPS from Gram-negative bacteria using TRPA1, TRPV1 and TLR4, mediating acute neurogenic inflammation, pain and sickness behavior ^8,16,17^. Additionally, they act as second sensory neurons along with respiratory epithelial cells such as pulmonary neuroendocrine cells (PNECs), brush cells, solitary chemoreceptor cells, and taste bud cells, collectively identifying pathogens ^1,18–22^. These epithelial cells sense airway signals and transmit them to sensory neurons. PNECs, in particular, participate in lung recognition processes, including responses to hypoxia, hypercarbia, mechanical stretch, allergens, water and extracellular traps ^22–25^. It has been reported that PNECs can produce CGRP and γ-Aminobutyric acid (GABA) to regulate pulmonary immune responses^20,24,26^. Many studies have shown that PNECs receive projections from various vagal sensory neurons, including *P2ry1* and *Pvalb* ^1,27,28^. However, it remains unclear whether and how vagal sensory neurons (i.e., nodose αCGRP^+^ neurons) feedforward regulate CGRP production in PNECs and trigger lung inflammation.

The preganglionic synapse of GABAergic neurons is finely regulated by GABA receptor signaling ^29^. In this study, we hypothesize that αCGRP^+^ nodose sensory neurons couple with PNECs to form a synapse. In response to the pulmonary LPS challenge, PNECs are activated to produce CGRP and GABA. PNEC-derived CGRP can amplify lung inflammation by increasing the production of proinflammatory cytokine and neutrophil infiltration, and GABA can modulate preganglionic GABA receptors in αCGRP^+^ nodose sensory neurons to limit the release of αCGRP. Through this feedback loop, CGRP production in PNECs will be fine-tuned. Overall, we sought to elucidate the neural circuits between the nodose ganglia and the PNECs through which the inflammatory response in the lungs is regulated.

To address these issues, we employed transcriptome analysis, tissue clearance imaging, electroexcitability recording, gene and cell-specific knockout, and other advanced methods. We found that pulmonary LPS challenge increased vagal excitability and CGRP expression in NG sensory neurons via TRPA1 rather than TLR4. Deletion of *Myd88* in *Phox2b*^+^ nodose sensory neurons reduced LPS-triggered vagal excitability and lung inflammation. A large number of nodose sensory neurons were TRPA1^+^αCGRP^+^ neurons, which innervated PNECs and alveoli. Deletion of *Trpa1* or *Calca* in the vagal ganglia reduced CGRP expression in PNECs, the proliferation of PNECs, and lung inflammation. Ablation of PNECs reduced LPS-induced lung inflammation, and lung GABA feedback modulated CGRP expression in nodose sensory neurons. Thus, we have established a nodose ganglion-pulmonary neuroepithelial circuit that regulates the inflammatory response in the lungs.

## Results

### The sensory neurons in nodose ganglion via MYD88 sense intratracheal LPS and promote lung inflammation

The major sensory neurons projected to the lungs are vagus ganglion sensory neurons and dorsal root ganglion sensory neurons. To distinguish between NG and DRG sensory neurons, we used *Phox2b^Cre^*mice that exhibited specificity for NG. By mating *Phox2b^Cre^* mice with Cre-dependent *Rosa26^tdTomato^* mice, we found that the *Phox2b^tdTomato+^* signal was specifically detected in the vagal ganglia but absent in the DRG (Extended Data Fig. 1a and 1c). To track which neurons in the nodose ganglia project to the lungs and respond to the LPS challenge, we injected Fast Blue into the lungs, a fluorescent dye that can retrogradely travel from the lungs to the vagal ganglia (Extended Data Fig. 1a). We observed that large amount of Fastblue^+^ staining was present in the *Phox2b^tdTomato+^*NG (Extended Data Fig. 1a), and only a small amount displayed in the T2 DRG (Extended Data Fig. 1c). By flow cytometry analysis, we found that about 4% of vagal sensory neurons were Fastblue^+^ (Extended Data Fig. 1b and 1c). Consistent with previous findings, these results suggest that sensory neurons projecting to the lungs primarily originate from the NG^30^.

Vagal sensory neurons can sense pathogens, and more in-depth research has focused on LPS of gram-negative bacteria, a model that can induce acute lung injury. The vagus nerve can recognize LPS injected intraperitoneally, producing specific neural electrical signals ^31^. However, it is unclear whether the nerve endings of vagal sensory neurons can sense LPS in the lungs. To investigate whether lung LPS challenge would cause a rapid vagal nerve response, we developed a unique lung LPS delivery method to bypass the upper airway. The vagal excitability is recorded by electrodes placing on the cervical vagus nerve after delivering either PBS or LPS (Fig. 1a). We found that both LPS and PBS challenge could elicit an increase in vagal firing rate after delivery. Lung PBS fluid stimulation induced vagal excitation within 1 minute, whereas LPS stimulation triggered strong nerve firing that persisted throughout the recording period. The normalized vagal compound action potentials (CAPs, detected neural events) recorded in the LPS-challenged group were higher compared to PBS-challenged group (Fig. 1b and 1c).

**Fig. 1.**
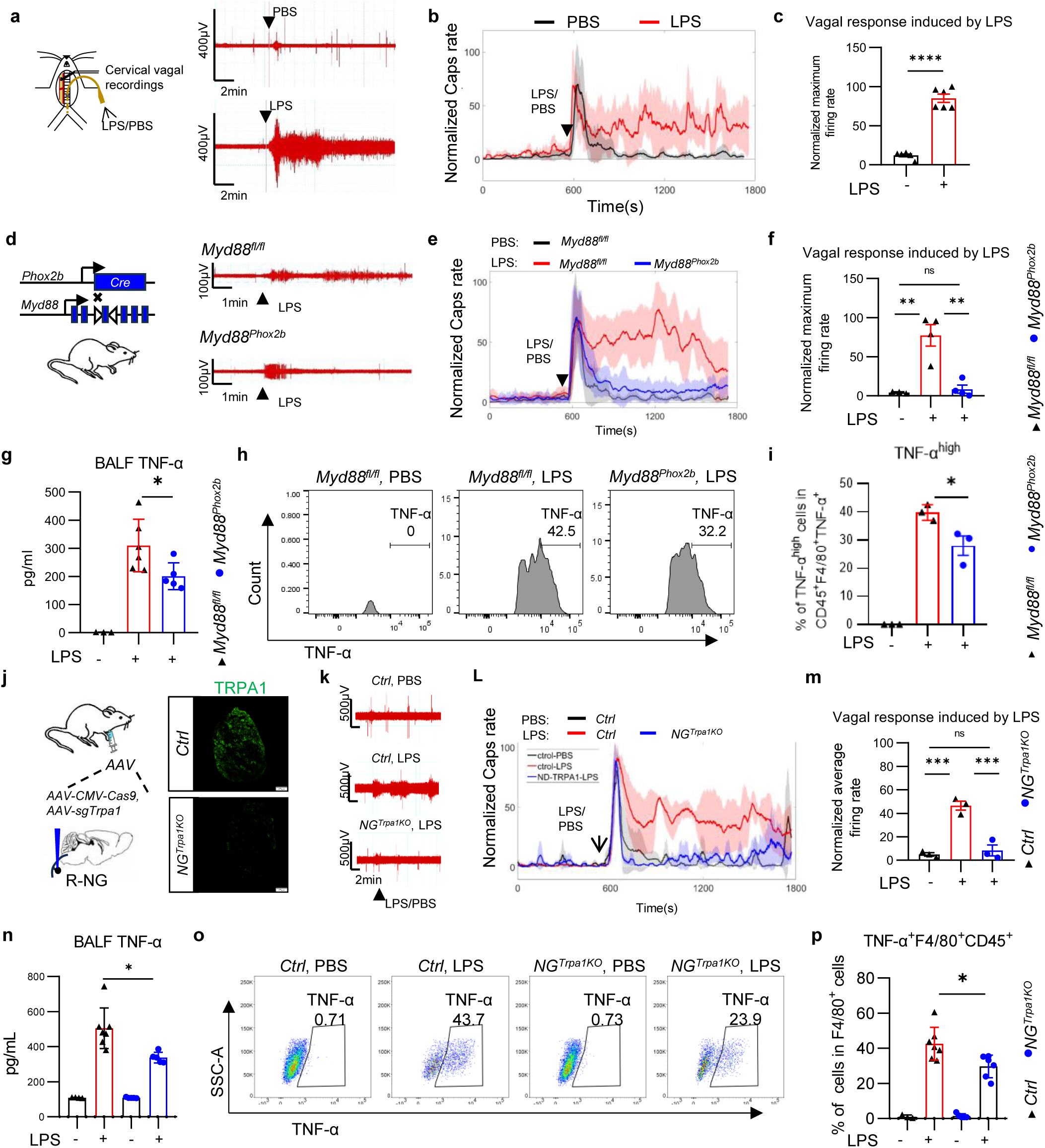
Nodose sensory neurons directly utilize TRPA1 to rapidly respond to intratracheal LPS and amplify lung inflammation. (a and b) In anesthetized wild-type mice, lung LPS stimulation was performed via endotracheal intubation, while recording the electrical activity of the cervical vagus nerve. Vagal responses to stimuli within the lung, including baseline (PBS, gray traces) and LPS (5 mg/kg, red traces) (n = 5-6 mice/group). (c) Quantification of peak responses to LPS (n = 5-6 mice/group). (d) Vagal responses to LPS stimuli within the lung in *Myd88^Phox2b^* mice and *Myd88^fl/fl^* mice. (n = 3-4 mice/group). (e) Vagal responses to baseline (PBS) and LPS (5mg/kg), including whether nodose neurons are Myd88 knockout (n = 3-4 mice/group). (f) Quantification of peak responses to LPS in *Myd88^Phox2b^* mice and *Myd88^fl/fl^* mice (n = 3-4 mice/group). (g) LPS induced a lower level of TNF-α in *Myd88^Phox2b^* mice compared with *Myd88^fl/fl^* mice in BALF (n = 3-6 mice/group). (h and i) LPS induced a lower level of TNFα in F4/80^+^ macrophages in the BALF of *Myd88^Phox2b^* mice compared with *Myd88^fl/fl^* mice (n = 3 mice/group). (j) Injection of *AAV-sgTrpa1* into the right nodose ganglion to generate *NG^Trpa1KO^* mice. (k and l) Vagal responses to LPS stimuli within the lung in *NG^Trpa1KO^* mice and Ctrl mice, including PBS for *Ctrl* mice (gray traces), LPS for *Ctrl* mice (red traces), and LPS for *Trpa1^KO^* mice (blue traces) (n = 5-7 mice/group). (m) Quantification of average responses to LPS in *NG^Trpa1KO^* mice and *Ctrl* mice (n = 3 mice/group). (n) LPS induced a lower level of TNF-α in *NG^Trpa1KO^* mice compared with *Ctrl* mice in BALF (n = 4-7 mice/group). (o and p) LPS induced a lower level of TNF-α in F4/80^+^ macrophages in the BALF of *NG^Trpa1KO^* mice compared with *Ctrl* mice (n = 4-7 mice/group). Scale bars: 100μm in (j). Student’s t test in (c). One-way ANOVA in (f), (g), (i), (m), (n) and (p). Mean ± SEM. ns, not significant. *p < 0.05, **p < 0.01, and ***p < 0.001.

MYD88 is a hub molecule of toll-like receptors that are required for pathogen sensing. It is broadly expressed in sensory neurons of the NG ^3^. We crossed *Myd88^fl/fl^*mice with *Phox2b^Cre^* mice to specifically delete *Myd88* in sensory neurons of the NG (*Myd88^Phox2b^* mice) (Fig. 1d). By recording the excitability of vagus nerve, we found that LPS-induced peak of Caps curve in *Myd88^fl/fl^*mice was abolished by deleting *Myd88* (Fig. 1e). Vagal nerve firing rate in LPS-challenged *Myd88^Phox2b^* mice was also reduced compared to LPS-challenged *Myd88^fl/fl^* mice (Fig. 1f). These data suggest that NG sensory neurons could sense lung LPS via MYD88 signaling to trigger vagus nerve firing.

To study the role of sensory neurons in mediating LPS-induced lung inflammation, we established an 1 h LPS challenged mouse model (Extended Data Fig. 2a). We found that lung LPS challenge at 1 h increased bronchoalveolar lavage fluid (BALF) TNF-α levels (Extended Data Fig. S2b), TNF-α expression in BALF F4/80^+^ cells (Extended Data Fig. 2c-2e), and lung polymorphonuclear neutrophil (PMN) infiltration (Extended Data Fig. 2f-2h). Using the above model, we found that TNF-α in BALF and TNF-α expression in F4/80^+^ macrophages were significantly reduced in LPS-challenged *MyD88 ^Phox2b^* mice compared to LPS-challenged *MyD88^fl/fl^* mice (Fig. 1g-1i). In LPS-challenged *MyD88 ^Phox2b^*, levels of LPS-induced *Il6, Cxcl1,* and *Ccl22* were also significantly reduced compared to the control group (Extended Data Fig. 3a and 3b). These findings indicate that sensory neurons of the NG via MYD88 sense LPS challenge to mediate lung inflammation.

### Pulmonary LPS-induced vagal responses rely on TRPA1^+^ rather than TLR4^+^ nodose sensory neurons

The TLR4 signaling pathway is a classical pathway for recognizing LPS. To clarify whether NG sensory neurons directly sense pulmonary LPS via TLR4, we recorded vagal excitability in *TLR4^KO^* and control mice after pulmonary LPS-challenge. Although TNF-α in BALF in LPS-challenged *TLR4^KO^* mice was significantly reduced compared to PBS-challenged control mice (Extended Data Fig. 4d), the normalized CAPs and firing rate in vagus nerve did not differ between pulmonary-LPS challenged *TLR4^KO^*and their counterparts (Extended Data Fig. 4a-4c). In addition, we developed an *in vivo* calcium imaging platform to record LPS-induced neural activity in the vagal nodose sensory neuron. We injected the calcium indicator AAV9-CMV-GCaMP6s into the vagal nodose ganglia, targeting GCaMP6s to all sensory neurons. Using a miniscope imaging system, we recorded the real-time responses of vagal neurons to LPS stimulation administered to the lungs. We found that LPS-induced calcium signaling in nodose ganglion sensory neurons persisted in TLR4 knockout mice (Extended Data Fig. 4i and 4j). These findings indicate that pulmonary LPS-induced vagal responses is not dependent on TLR4^+^ nodose sensory neurons.

In addition to TLR4, it is reported that LPS can directly activate TRPA1 and TRPV1 expressed in neurons ^16,17^. By reanalyzing the single-cell RNA sequencing datasets from Chang’s group^32^, we found that *Trpa1* and *Myd88* genes are highly expressed in *Prdm12^+^* jugular and *Phox2b^+^* nodose sensory neurons (Extended Data Fig. 4e-4f). However, *Tlr4^+^* sensory neurons account for 10.6% in the jugular ganglia and only 1.8% in the nodose ganglia (Extended Data Fig. 4g). Strikingly, 53% of the *Trpa1*^+^ NG neurons projected to the lungs were much higher than the 1.8% of *Tlr4*^+^ neurons (Extended Data Fig. 4h). *Trpa1*^+^ lung-innervated neurons express lower levels of *Tlr4*, suggesting that TRPA1 is more likely to function as a sensor for LPS in the pulmonary physiological environment.

In QZ2 labeled NG sensory neurons projected to the lungs*, Trpa1^+^* sensory neurons expressed *Myd88*^32^, a hub molecule used for LPS sensing (Extended Data Fig. 5a). We next investigated whether activation of TRPA1 enhances MyD88 oligomerization to facilitate signal transduction, we constructed Neuro-2a cells expressing *MyD88-GFP (Neuro-2a-MyD88-GFP)* (Extended Data Fig. 5b), and overexpressed TRPA1 in Neuro-2a-MyD88-GFP cells. In Neuro-2a-MyD88-GFP cells overexpressing TRPA1, the number and size of MyD88-GFP puncta were significantly increased (Extended Data Fig. 5c and 5d). This indicates that LPS can activate TRPA1 in neurons to enhance MyD88 oligomerization for intracellular signal transduction.

We evaluated the in vivo impact of TRPA1 on vagal signaling induced by LPS by selectively knocking out TRPA1 in the NG. We injected *AAV-CMV-Cas9* and *AAV-sgTrpa1* adenovirus targeting TRPA1 expression into the right nodose ganglion of mice (*NG^Trpa1KO^*mice), knocking out *Trpa1* expression within the nodose ganglion (Fig. 1j). In *NG^Trpa1KO^* mice, the vagal response to the lung LPS stimulation was significantly reduced (Fig. 1k-1m). Regarding the regulation of pulmonary inflammation, compared to the control group, *NG^Trpa1KO^* mice showed significantly reduced TNF-α expression in BALF (Fig. 1o). Furthermore, there was a significant decrease in the expression level of TNF in F4/80 cells in the BALF (Fig. 1p and 1q). Taken together, these results suggest that nodose sensory neurons can utilize TRPA1 to sense and respond to pulmonary LPS within the pulmonary physiological environment, upregulating CGRP expression and promoting pulmonary inflammation responses.

### NG sensory neurons upregulate *αCGRP* expression in response to Lung LPS challenge

To study transcriptome changes in NG sensory neurons in response to lung LPS challenge, we isolated vagal ganglia to perform bulk RNAseq analysis (Fig. 2a and Extended Data Fig. 6a). We identified 886 differentially expressed genes, among which 463 were significantly upregulated and 423 were significantly downregulated (Extended Data Fig. 6b). We visualized these differentially expressed genes in a heatmap and performed GO-biological process (GO-BP) enrichment analysis (Extended Data Fig. 6c and 6d). The enriched pathways included regulation of nervous system process, detection of external stimulus, regulation of sensory perception, detection of temperature stimulus, and sensory perception of pain (Extended Data Fig. 6d). These pathways suggest that the vagal sensory neurons recognize and respond to lung LPS challenge. Additionally, the differentially expressed genes were enriched in regulation of epithelial cell proliferation (Extended Data Fig. 6d), suggesting that the vagal sensory neurons might exert neuro-immune regulatory effects by modulating lung epithelial cells in response to lung LPS.

**Fig. 2.**
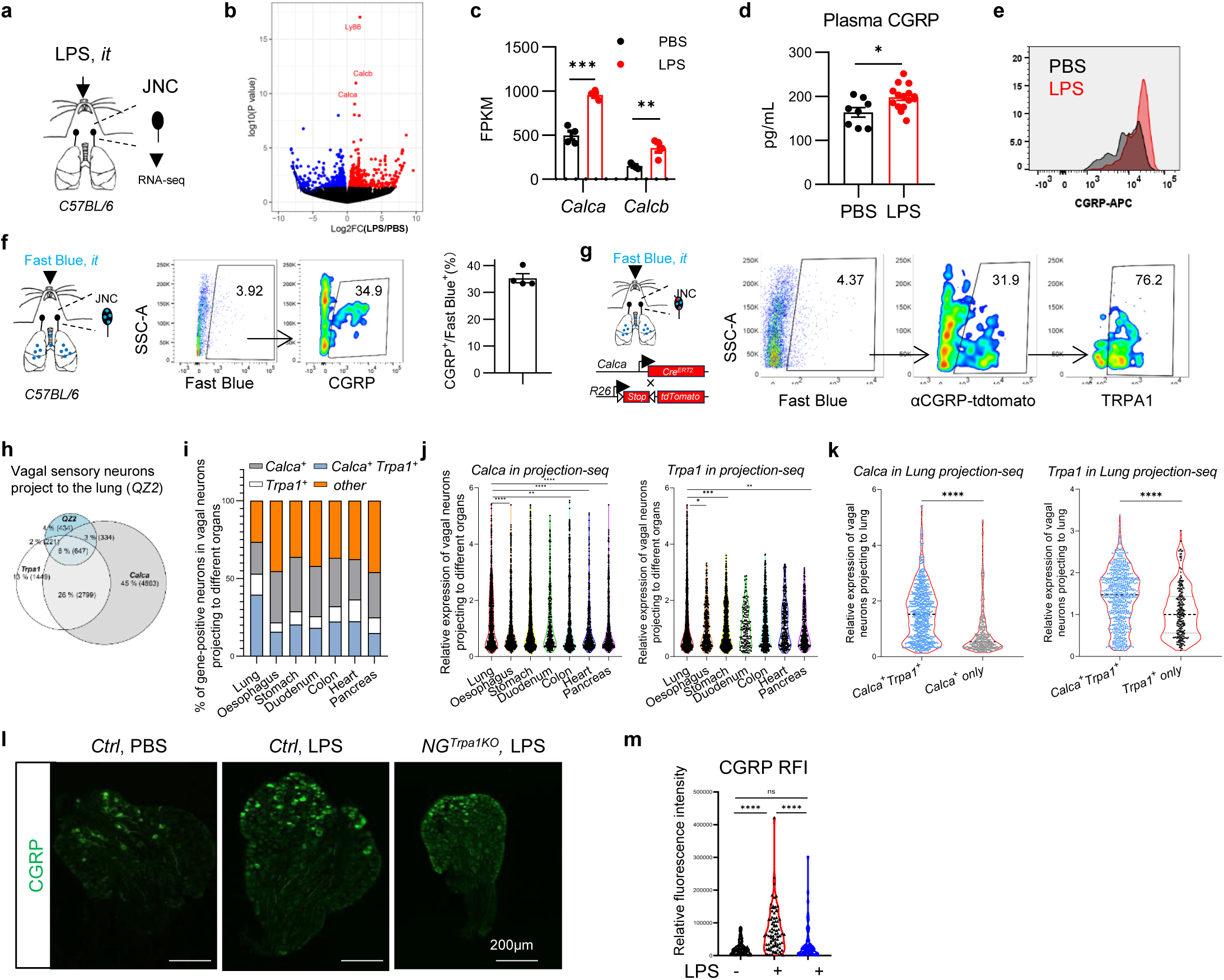
αCGRP^+^Trpa1^+^ nodose sensory neurons projecting to the lungs respond to pulmonary LPS stimulation. **(A)** (a) Schematic diagram of RNA-seq analysis using isolated nodose ganglia following pulmonary LPS administration. (b) RNA-seq volcano plot of vagal ganglion treated or not treated with LPS (5 mg/kg) for 1 h. The neurotransmitter-related and immune-related genes are labeled. (n = 4 mice/group). (c) Analysis of CGRP gene expression in vagal ganglion upon LPS treatment via RNA-seq. (n = 4 mice/group). (d) LPS induced a larger increase of CGRP levels in plasma (n = 8-16 mice/group). (e) Flow cytometric analysis of CGRP expression intensity in the vagal ganglion. (f) The proportion of CGRP-positive sensory neurons among all vagal sensory neurons projecting to the lung was determined using Fast Blue retrograde tracing. (n = 4 mice/group). (g) The proportion of TRPA1-positive sensory neurons among vagal αCGRP-positive sensory neurons projecting to the lung in *Calca^tdTomato^* mice. (h) The expression of *Calca* and *Trpa1* genes in QZ2^+^ neurons, which are vagal neurons projecting to the lungs^32^. (i) The distribution of vagal *Trpa1/Calca* double-positive neurons, *Trpa1*-positive neurons, and *Calca*-positive neurons projecting to the lungs and other locations. (j) The expression levels of *Trpa1* and *Calca* in neurons projecting to the lungs and other locations. (k) The expression levels of *Trpa1* and *Calca* in *Trpa1*/*Calca* double-positive neurons, *Trpa1*^+^ neurons, and *Calca*^+^ neurons that project to the lungs. (l and m) CGRP mean fluorescence intensity in the nodose ganglion of *NG^Trpa1KO^*mice and *Ctrl* mice. (n = 4– 7 mice/group). Scale bars: 200 μm in (l). Student’s t test in (c), (d), (j) and (k). One-way ANOVA in (m). Mean ± SEM. ns, not significant. *p < 0.05, **p < 0.01, and ***p < 0.001.

Using volcano plot analysis, we found that the neuropeptide genes *Calca* (encoding α-CGRP) and *Calcb* (encoding β-CGRP) were among the top differentially upregulated genes in the vagal ganglia from LPS-challenged mice (Fig. 2b and 2c). Consistently, CGRP levels in the plasma from the LPS-challenged mice were also increased compared to the control (Fig. 2d). By flow cytometry and tissue clearing imaging analyses, we found that CGRP expression was higher in the nodose sensory neurons from the LPS challenged group (Fig. 2e). Given that the expression level of *Calca* is higher than that of *Calcb*, and the number of *Calca^+^* nodose sensory neurons projecting to the lungs is significantly greater than that of *Calcb^+^* neurons, we focused our analysis on αCGRP-positive sensory neurons (Fig. 2c and Extended Data Fig. 4e). Furthermore, the differentially expressed gene *Calca* was enriched in the GO-BP pathways detection of temperature stimulus, feeding behavior, sensory perception of pain, detection of abiotic stimulus, and detection of external stimulus, with four of these pathways belonging to detection of stimulus (Extended Data Fig. 6d).

By GO analysis, we found that differential genes related to “Response to stimulus,” “Nervous system,” “Axon,” and “Synapse,” were enriched (Extended Data Fig. 7a and 7b). We further analyzed synaptic pathways and found differential genes regarding “Regulation of dopamine secretion”, “Dopamine receptor signaling pathway”, “Sensory perception”, “Response to stimulus”, “GABAergic synapse”, and “Glutamatergic synapse” were prominently changed (Extended Data Fig. 7c and 7d). These findings indicate that lung LPS challenge could stimulate synapse formation in different groups of sensory neurons in vagal ganglia. We further conducted dimensionality reduction and unsupervised classification to dissect lung LPS-triggered vagus nerve electrical signals. We found that five clusters of responders could consistently respond to the pulmonary LPS challenge and characterize it with special waveforms (Extended Data Fig. 7e-7g). These findings suggest that different clusters of vagal sensory neurons are involved in sensing and responding to pulmonary LPS challenge.

Based on the data showing *Trpa1^+^* nodose sensory neurons responding to LPS, we hypothesize that activation of nodose *Trpa1^+^* sensory neurons in the nodose ganglion will upregulate CGRP to respond to LPS-induced lung inflammation. To test this hypothesis, we first confirmed the projection of CGRP^+^ sensory neurons to the lung. By isolating the vagal ganglia from mice that received an intratracheal injection of Fast Blue, we found that 3.92% of neurons in the vagal ganglia are fast blue^+^ and thus project to the lung, while 34.9% of neurons within the fast blue^+^ population were CGRP^+^ neurons (Fig. 2f). To further clarify the co-expression of αCGRP and TRPA1 in sensory neurons projecting to the lung, we crossed *Calca^CreERT2^* mice with Cre-dependent Rosa26-tdTomato mice (*αCGRP^tdTomato^* mice) to specifically label αCGRP-expressing neurons projecting to the lung. Through neuroanatomical tracing, we observed that approximately 76.2% of αCGRP^+^ sensory neurons projecting to the lungs were TRPA1-positive neurons (Fig. 2g).

To elucidate the characteristics of nodose *αCGRP^+^Trpa1^+^* neurons, we reanalyzed the projection-seq data of vagal neurons conducted by Chang and colleagues^32^. We found distinct expression patterns of *Trpa1^+^* and *Calca*^+^ in vagal neurons projecting to different visceral organs (Fig. 2h and Extended Data Fig. 9h). By scRNAseq analysis^32^, QZ2 labeled NG sensory neurons projected to the lungs, the composition of these sensory neurons was: *Trpa1^+^Calca^+^* (39.5%), other (26.5%), *Calca^+^* (20.4%), and *Trpa1^+^ (*13.5%) (Extended Data Fig. 4e and 4f). Specifically, 39.5% of sensory neurons projecting to the lungs were *Trpa1^+^Calca^+^* double-positive, which was significantly higher compared to other visceral tissues including the esophagus, stomach, duodenum, colon, heart, and pancreas (Fig. 5i). Not only was the proportion higher in lung-projecting neurons, but the expression levels of *Trpa1* and *Calca* were also elevated compared to other tissues (Fig. 5j). Further analysis of sensory neurons projecting to the lungs revealed significantly higher expression levels of *Calca* and *Trpa1* in double-positive neurons compared to single-positive neurons (Fig. 5k). This suggests that nodose *Trpa1^+^Calca^+^*sensory neurons are more likely to sense LPS and release αCGRP to regulate LPS-induced pulmonary inflammation.

To validate the direct role of TRPA1 in mediating CGRP production, we used mice with TRPA1 knocked out in the NG (*NG^Trpa1KO^* mice) and observed an increase in CGRP expression within the NG induced by LPS stimulation. However, we did not detect a rise in CGRP expression within the NG in *NG^Trpa1KO^* mice following LPS administration (Fig. 2l and 2m).

### αCGRP^+^ nodose sensory neurons densely innervate NEBs

In the lungs, nodose sensory neurons are primarily distributed beneath the large airways (Extended Data Fig. 1d), projecting to the airway bifurcations where they converge into neuroepithelial bodies (NEBs) composed of PNECs. We examined the distribution of αCGRP sensory neurons in the lungs of *αCGRP^tdTomato^*mice and observed that αCGRP^+^ sensory neurons densely projected to PNECs (Fig. 3a). To confirm the contribution of nodose ganglia, we crossed *Phoxb^Cre^* mice with Cre-dependent *Rosa26-tdTomato* mice and co-labeled with CGRP to mark nodose sensory neurons in the lungs. While some nodose sensory neurons project into the interior of NEBs, CGRP-positive nodose ganglion sensory neurons are located beneath NEBs and do not penetrate into the NEB interiors (Fig. 2b and 2c). We conducted a statistical analysis of the nerve fibers surrounding the NEBs and observed that approximately 40% of sensory neurons projected to the NEBs were CGRP^+^ neurons (Fig. 3d). Additionally, we observed that CGRP^+^ nodose sensory neurons projected to the alveoli, accounting for 41% of all nodose sensory neurons projecting to alveoli (Fig. 3e and 3f).

**Fig. 3.**
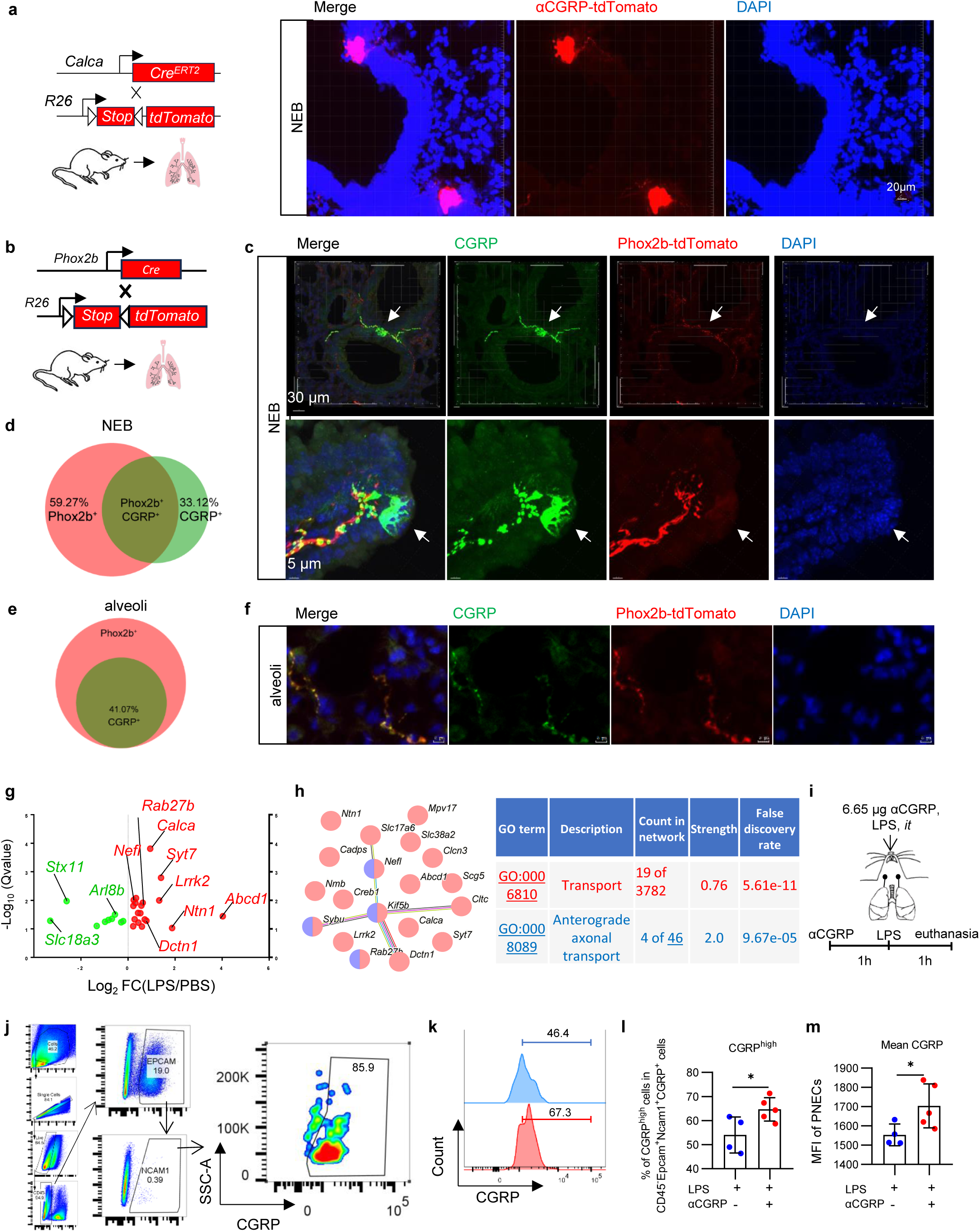
αCGRP^+^Trpa1^+^ nodose sensory neurons innervate PNECs, promoting excessive CGRP production in PNECs. (a) In *Calca^tdTomato^* mice, detect the projection of CGRP-positive neurons onto PNEC cells. (b and c) In *Phox2b^tdTomato^*mice, two-color immunohistochemistry for CGRP (green) and Phox2b-tdTomato (red) was performed to label CGRP-positive nodose neurons and PNECs at airway bifurcations. (d) The proportion of CGRP-positive sensory neurons among the nodose sensory neurons projecting to NEBs. (e) In *Phox2b^tdTomato^* mice, two-color immunohistochemistry for CGRP (green) and Phox2b-tdTomato (red) was conducted to identify CGRP-positive nodose neurons within the alveolar region. (f) Vagal sensory neuronal transport under LPS stimulation was analyzed using RNAseq databases of the vagal ganglion. (h) 19 transport genes in vagal ganglia, along with their enriched GO pathways, were induced by pulmonary LPS stimulation. (i) Administration of LPS or αCGRP+LPS into the lungs of mice. (j) Gating strategies for CGRP^+^EpCAM^+^NCAM^+^CD45^−^ PNEC used in flow cytometry. (k and l) The proportion of highly CGRP-expressing PNECs among all CGRP^+^EPCAM^+^NCAM^+^ cells in lung tissue from αCGRP-LPS-treated mice compared to LPS-treated mice. (m) Comparison of CGRP mean fluorescence intensity in CGRP^+^EpCAM^+^NCAM^+^ cells in lung tissue from αCGRP-LPS-treated mice compared to LPS-treated mice. Scale bars: 30 μm in the upper part of (b). 20 μm in (a). 5 μm in the lower part of (b) and (f). Student’s t test in (l) and (m). One-way ANOVA in (k). Mean ± SEM. ns, not significant. *p < 0.05, **p < 0.01, and ***p < 0.001.

### LPS challenge enhances vagal axonal transport and promotes CGRP-induced CGRP expression in PNECs

To test whether LPS challenge affects vagal sensory neuronal transport, we analyzed the vagal ganglion RNAseq databases. We found that LPS challenged in the mouse lungs increased 19 transport genes in vagal ganglia. Among the 19 genes, *Rab27b*, *Sybu*, *Nefl*, and *Kif5b* were associated with anterograde axonal transport. *Syt7* (synaptotagmin 7) and *Lrrk2* have been reported to mediate neuropeptide release or secretion in neurons^33,34^ (Fig. 3g-3c).

The activation of αCGRP^+^ sensory neurons and the release of αCGRP not only function as a neurotransmitter but also act as a neuroimmune regulatory substance, participating in various immune and inflammatory responses, particularly in the immune reactions of the lungs and airways^4,24,26^. PNECs are specialized epithelial cells that are the primary cellular source of CGRP in the lungs, expressing both the epithelial marker EpCAM and the neural adhesion molecule NCAM1 (Extended Data Fig. 8a) ^1^. We analyzed CGRP^+^EpCAM^+^NCAM1^+^CD45^−^ cells by flow cytometry to identify PNEC cells in the lungs. The results showed that approximately 80∼90% of CGRP^+^ lung cells were EpCAM^+^NCAM1^+^CD45^−^ (Fig. 3j). To investigate whether αCGRP regulates CGRP within PNECs, we intratracheally instilled αCGRP into mouse lungs, followed by LPS administration (Fig. 3i). In CGRP + LPS treated mice, CGRP^high^EpCAM^+^NCAM1^+^ cells and intensity of CGRP expression in CGRP^+^EpCAM^+^NCAM1^+^ cells were increased compared to LPS-challenged mice (Fig. 3k-3m). Meanwhile, we used a cell model to investigate whether LPS can enhance αCGRP-induced CGRP expression in PNECs. To this end, we treated the SCLC #1 cell line (a model of PNECs^35^) with αCGRP and LPS. Consistent with *in vivo* experiments, we found that both extracellular and intracellular CGRP were increased in LPS + αCGRP-treated PNECs compared to the LPS-treated group (Extended Data Fig. 8b-8d). CGRP activation of its receptors is known to increase cAMP/PKA signaling ^36^, which is also a key signal to enhance CGRP expression^37^. PNEC cell line treated with the PKA inhibitor H-89 significantly reduced the levels of CGRP in the cell supernatant without αCGRP treatment (Extended Data Fig. 8e and 8h). However, in the presence of αCGRP, H-89 surprisingly increased CGRP levels in the supernatant and cell lysate compared to the αCGRP-treated group (Extended Data Fig. 8e and 8h). Then, we silenced *Prkaca* and *Prkacb* expression in PNEC cells using lentivirus-mediated specific shRNA gene silencing to investigate whether PKA regulates CGRP synthesis in PNECs. We found that knockdown of *Prkaca* and *Prkacb* in PNEC cells increased CGRP gene (*Calca*, *Calcb*) expression levels compared to the control group (Extended Data Fig. 8i and 8k). Additionally, under αCGRP treatment, knockdown of *Prkaca* and *Prkacb* in PNEC cells also increased CGRP protein expression levels compared to the control group (Extended Data Fig. 8f and 8h). These findings suggest that the role of PKA in regulating CGRP expression in PNECs is different from the results of previous studies^38–40^.

### Nodose TRPA1^+^αCGRP^+^ sensory neurons induce CGRP expression in PNECs and amplify lung inflammation

Next, we tested whether αCGRP-positive nodose sensory neurons regulate CGRP expression in PNECs. We injected *Calca^Cre^Rosa26^DTR^* mice with diphtheria toxin (DT) into NG to ablate CGRP neurons (named *NG^CalcaABLATE^* mice). The *control* group were age-matched Cre-negative littermates received DT. (Fig. 4a). We found that ablation of CGRP neurons within the NG relatively preserved CGRP neurons in JG (Fig. 4b) and did not alter the quantity of PNECs and NEB structure in the lungs of *NG^CalcaABLATE^* mice (Fig. 4c). However, flow cytometric analysis revealed that both CGRP^high^ cells and CGRP intensity in lung CGRP^+^EpCAM^+^NCAM1^+^ population were reduced in LPS-challenged *NG^CalcaABLATE^* mice compared to the LPS-treated control mice (Fig. 4d-F). The lung CD45^−^ EpCAM^+^NCAM1^+^ cells were decreased after LPS injury (Fig. 5G) and TNF expression in BALF F4/80^+^ macrophages was also reduced in LPS-challenged *NG^CalcaABLATE^*mice compared to LPS-treated control group (Fig. 4g-4i).

**Fig. 4.**
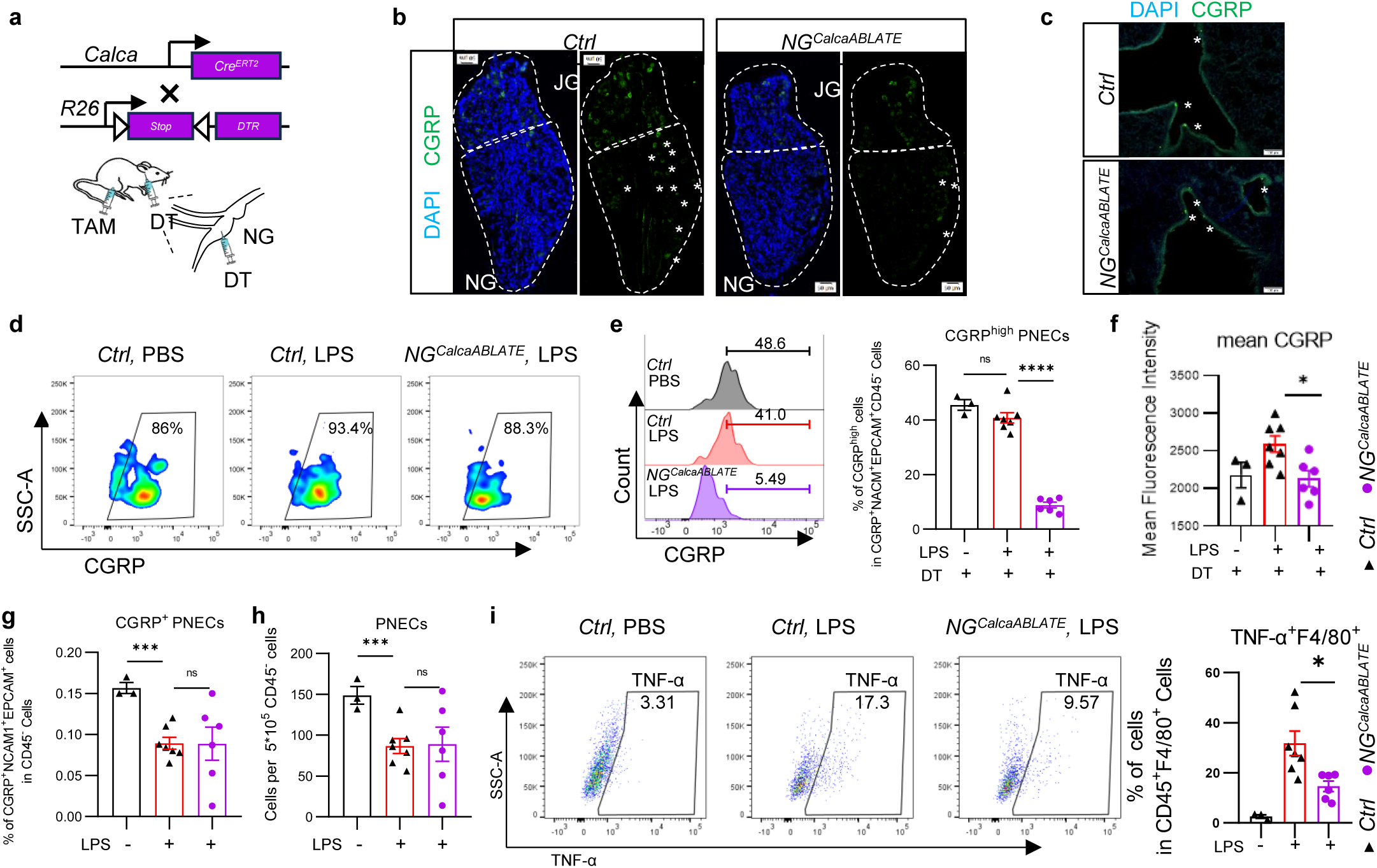
Nodose αCGRP^+^ sensory neurons deficiency results in reduced CGRP expression levels in PNECs and reduced lung inflammation. (a) Diphtheria Toxin (DT) was injected into the nodose ganglia in *Calca^CreERT2^; loxP-DTR* (*NG^CalcaABLATE^*) mice and *loxP-DTR* (*Ctrl*) mice. (b and c) CGRP immunostaining (green) in the nodose/jugular complex (B) and lung (C) of *Calca^CreERT2^; loxP-DTR* (*NG^CalcaABLATE^*) mice and *loxP-DTR* (*Ctrl*) mice following NG injection, with DAPI nuclear counterstaining. (d-e) The gating strategy of CGRP^+^EPCAM^+^NCAM^+^ PNECs from *NG^CalcaABLATE^* mice and *Ctrl* mice following LPS stimulation (D). The proportion of highly CGRP-expressing PNECs among all CGRP^+^EPCAM^+^NCAM^+^ cells in lung tissue from *NG^CalcaABLATE^* mice and *Ctrl* mice following LPS stimulation (E). (n = 3-7 mice/group) (f) Comparison of CGRP mean fluorescence intensity in CGRP^+^EpCAM^+^NCAM^+^ cells in lung tissue from *NG^CalcaABLATE^* mice and *Ctrl* mice following LPS stimulation. (n = 3-7 mice/group) (g) The proportion of CGRP^+^EPCAM^+^NCAM^+^ cells among all CD45^−^ cells in lung tissue from *NG^CalcaABLATE^* mice and *Ctrl* mice following LPS stimulation. (n = 3-7 mice/group) (h) The number of PNECs per 5*10^5^ CD45^−^ cells in lung tissue from *NG^CalcaABLATE^* mice and *Ctrl* mice following LPS stimulation. (n = 3-7 mice/group) (i) LPS induced a lower level of TNFα in F4/80^+^ macrophages in the BALF of *NG^CalcaABLATE^* mice compared with *Ctrl* mice (n = 3-7 mice/group). Scale bars: 100 μm in (c). 50 μm in (b). One-way ANOVA in (e), (f), (g), (h) and (i). Mean ± SEM. ns, not significant. *p < 0.05, **p < 0.01, and ***p < 0.001.

**Fig. 5.**
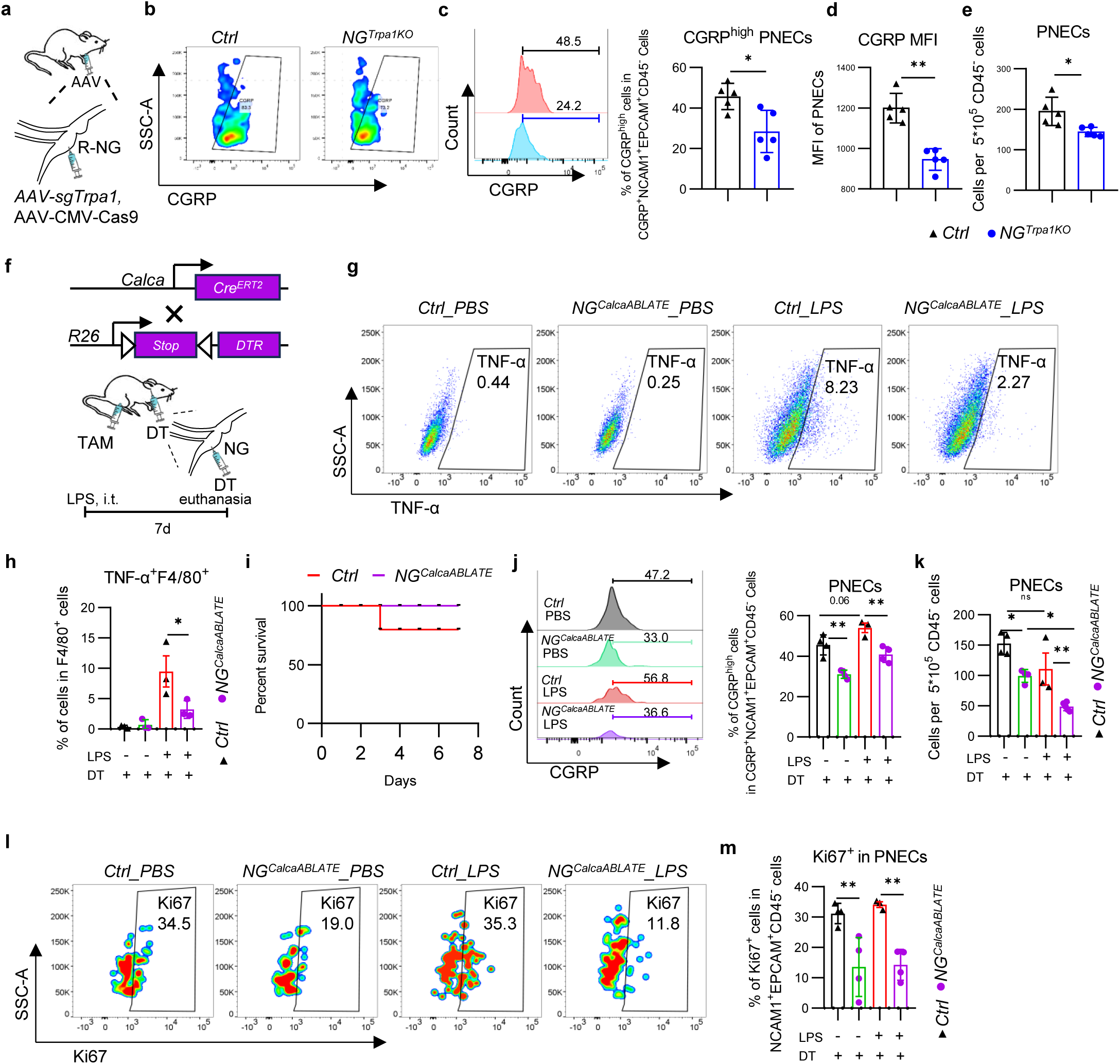
Nodose Trpa1^+^αCGRP^+^ sensory neurons are necessary for PNECs proliferation. (a) The injections of *AAV-sgTrpa1* and *AAV-sgScramble* into the right nodose ganglion are performed to generate *NG^Trpa1KO^*mice and *Ctrl* mice, respectively. (b) The gating strategy of CGRP^+^EPCAM^+^NCAM^+^ PNECs from *NG^Trpa1KO^* mice and *Ctrl* mice. (c) The proportion of highly CGRP-expressing PNECs among all CGRP^+^EPCAM^+^NCAM^+^ cells in lung tissue from *NG^Trpa1KO^* mice and *Ctrl* mice (n = 5 mice/group). (d) Comparison of CGRP mean fluorescence intensity in CGRP^+^EpCAM^+^NCAM^+^ cells in lung tissue from *NG^Trpa1KO^* mice and *Ctrl* mice (n = 5 mice/group). (e) The number of PNECs per 5*10^5^ CD45^−^ cells in lung tissue from *NG^Trpa1KO^*mice and *Ctrl* mice following LPS stimulation (n = 5 mice/group). (f) Diphtheria Toxin (DT) was injected into the nodose ganglia in *Calca^CreERT2^; loxP-DTR* (*NG^CalcaABLATE^*) mice and *loxP-DTR* (*Ctrl*) mice. Three weeks after DT injection, a 7-day LPS stimulation model was established. (g and h) During the 7 days of LPS stimulation, *NG^CalcaABLATE^* mice exhibited lower levels of TNF-α in BALF compared to *Ctrl* mice (n = 3 mice/group). (i) The mortality rate in *NG^CalcaABLATE^* mice and *Ctrl* mice during 7 days of LPS stimulation. (j) The proportion of highly CGRP-expressing PNECs among all CGRP^+^EPCAM^+^NCAM^+^ cells in lung tissue from *NG^CalcaABLATE^* mice and *Ctrl* mice following 7 days of LPS stimulation (n = 3-4 mice/group). (k) The number of PNECs per 5*10^5^ CD45^−^ cells in lung tissue from *NG^CalcaABLATE^* mice and *Ctrl* mice following 7 days of LPS stimulation (n = 3-4 mice/group). (l and m) The proportion of Ki67^+^ PNECs among all CGRP^+^EPCAM^+^NCAM^+^ cells in lung tissue from *NG^CalcaABLATE^* mice and *Ctrl* mice following 7 days of LPS stimulation (n = 3-4 mice/group). Student’s t test in (c), (d) and (e). One-way ANOVA in (h), (j), (k) and (m). Mean ± SEM. ns, not significant. *p < 0.05, **p < 0.01, and ***p < 0.001.

We repeated the experiments in *NG^Trpa1KO^* mice (receiving vector targeting deletion of *Trpa1* in vagal ganglia) (Fig. 5a) and found that both CGRP^+^ cells and CGRP intensity in lung CGRP^+^EpCAM^+^NCAM1^+^ population from LPS-challenged *NG^Trpa1KO^*mice were decreased compared with LPS-challenged *control* mice (Fig. 5b-5d). These findings suggest that TRPA1^+^ neurons in NG innervate PNECs and regulate their CGRP production under LPS challenge. In collaboration with our findings, *TRPA1^+^αCGRP^+^*neurons in NG rather than the other cell populations are most likely to release αCGRP that could boost CGRP expression in PNECs under lung LPS challenge.

### Nodose TRPA1^+^αCGRP^+^ sensory neurons regulate the proliferation of PNECs

In the RNA-seq database of the nodose ganglion induced by LPS, differential pathway analysis revealed enrichment in “regulation of epithelial cell proliferation.” This suggests that nodose TRPA1^+^αCGRP^+^ sensory neurons may be involved in regulating the proliferation of PNECs. In *NG^Trpa1KO^* mice, which have a selective knockout of TRPA1 in the nodose ganglion, we found that the number of CGRP^+^EpCAM^+^NCAM1^+^ PNECs was significantly lower compared to *control* mice (Fig. 5e). To investigate the effect of αCGRP released by the activation of nodose TRPA1^+^αCGRP^+^ sensory neurons on PNEC proliferation, we used *NG^CalcaABLATE^*mice, which selectively ablate αCGRP^+^ sensory neurons in the nodose ganglion, to establish a 7-day LPS treatment animal model (Fig. 5f). Consistent with the 1-hour LPS stimulation model, the TNF-α levels in the BALF of *NG^CalcaABLATE^* mice were significantly lower than those in *control* mice (Fig. 5g and 5h). Furthermore, by recording the LPS-induced mortality, we found that selective ablation of αCGRP^+^ sensory neurons in the nodose ganglion reduced the mortality of the mice (Fig. 5i). Regarding CGRP levels in PNECs, selective ablation of αCGRP^+^ sensory neurons significantly reduced the CGRP levels in PNECs (Fig. 5j). Additionally, the number of PNECs was significantly reduced following selective ablation of αCGRP^+^ sensory neurons, and PNEC regeneration was significantly impaired compared to *control* mice (Fig. 5k). We assessed cell proliferation ability using Ki67 staining and found that the Ki67 levels in the CGRP^+^EpCAM^+^NCAM1^+^ cell population were significantly reduced in *NG^CalcaABLATE^* mice compared to *control* mice (Fig. 5l and 5m). These data suggest that the activation of TRPA1^+^αCGRP^+^ nodose sensory neurons and the release of αCGRP not only regulate the CGRP levels within PNECs but also modulate the number of PNECs, while participating in the regulation of LPS-induced pulmonary inflammation.

### PNECs and CGRP amplify LPS-induced lung inflammation

To examine whether PNECs would contribute to LPS-induced lung inflammation, we developed *Ascl1^ABLATE^* mice (*Ascl1^CreERT2^* mice crossed with Cre-dependent *Rosa26-DTR* mice and then received an intraperitoneal injection of DT to ablate PNECs) (Fig. 6a). In these mice, CGRP^+^ NEBs were depleted confirmed by immunofluorescence (Fig. 6a). The population of lung EpCAM^+^NCAM1^+^ cells were also reduced compared to the control group (Fig. 6c). We recorded the vagal nerve responses to lung LPS in *Ascl1^ABLATE^* mice and *Control* mice and observed that the ablation of PNECs does not affect the vagal nerve’s electrical activity in response to LPS. This further confirms our previous findings that the vagal nerve utilizes TRPA1 to directly sense and respond to lung LPS (Fig. 6d and 6e). Furthermore, under LPS challenge, *Ascl1^ABLATE^* mice showed less severe lung inflammation evidenced by a significant reduction in BALF TNF-α and TNF-α^+^F480^+^ macrophages (Fig. 6F-6h). Considering that polymorphonuclear neutrophils (PMNs) plays a central role in lung inflammation^41^, we performed flow cytometry and found that Ly6G^high^Ly6C^low^CD11b^+^ PMNs were reduced in LPS-challenged *Ascl1^ABLATE^* mice compared to their controls (Fig. 6i and 6j). The lung inflammatory genes *il6* and *Ccl2* were also decreased in LPS-treated *Ascl1^ABLATE^* mice (Fig. 6k). We also monitored the LPS-induced mortality in *Ascl1^ABLATE^* mice and control mice, and found that ablation of PNECs significantly reduced LPS-induced mortality (Fig. 6i).

**Fig. 6.**
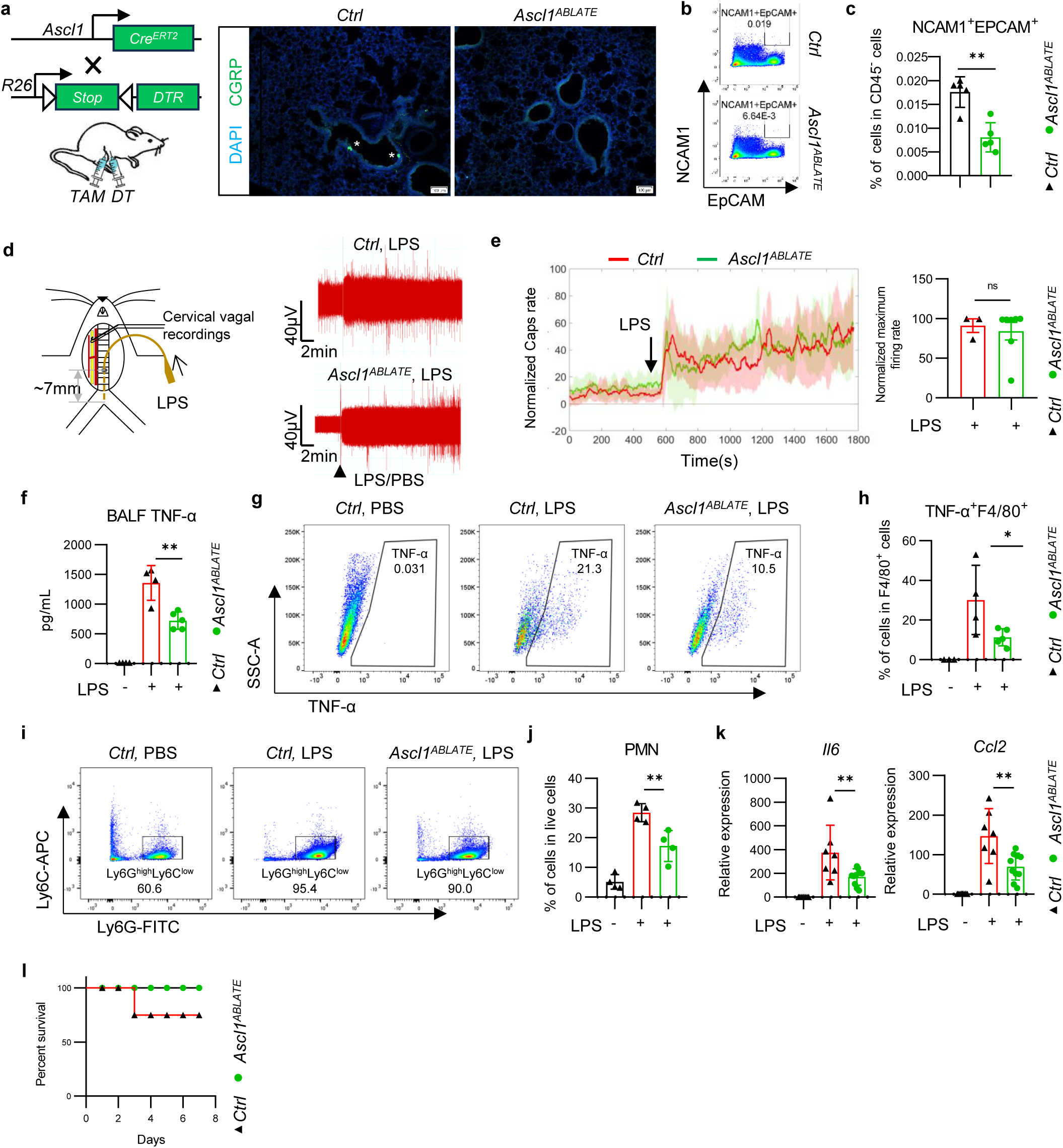
PNECs deficiency alleviated LPS-induced pulmonary inflammation. (a) Sequential intraperitoneal injections of tamoxifen and diphtheria toxin were used to genetically ablate PNECs in Ascl1CreERT2; loxP-DTR (*Ascl1^ABLATE^*) mice and loxP-DTR (*Ctrl*) mice. (b and c) The proportion of EPCAM^+^NCAM^+^ cells among CD45-cells was assessed in *Ascl1^ABLATE^* mice and *Ctrl* mice. (n = 5 mice/group). (d) Vagal responses to LPS stimuli within the lung in *Ascl1^ABLATE^* mice and *Ctrl* mice. (n = 3-7 mice/group) (e) Quantification of maximum responses to LPS in Trpa1KO mice and Ctrl mice (n = 3 mice/group). (f) Following LPS stimulation, the level of TNF-α in the BALF of *Ascl1^ABLATE^* mice was lower compared to that in *Ctrl* mice (n =4-5 mice/group). (g and h) Following LPS stimulation, *Ascl1^ABLATE^* mice exhibited a lower level of TNF-α in F4/80^+^ macrophages in the BALF compared to *Ctrl* mice (n = 4-5 mice/group). (i) The proportion of Ly6G^high^Ly6C^low^ cells among CD11B^+^ cells in lung tissue was assessed in *Ascl1^ABLATE^* mice and *Ctrl* mice following LPS stimulation (n = 4-5 mice/group). (j) The proportion of Ly6G^high^Ly6C^low^CD11b^+^ cells (PMNs) among live cells in lung tissue was assessed in *Ascl1^ABLATE^* mice and *Ctrl* mice following LPS stimulation (n = 4 mice/group). (k) *Il6* and *Ccl2* expression levels in lung tissue were assessed in *Ascl1^ABLATE^* mice and *Ctrl* mice following LPS stimulation. (n = 6-10 mice/group) (l) The mortality rate in *Ascl1^ABLATE^* mice and *Ctrl* mice during 7 days of LPS stimulation. Scale bars:100 μm in (a). Student’s t test in (c) and (e).One-way ANOVA in (f), (h), (j) and (k). Mean ± SEM. ns, not significant. *p < 0.05, **p < 0.01, and ***p < 0.001.

To test the effect of αCGRP on LPS-elicited lung inflammation, we intratracheally instilled CGRP into mouse lungs, followed by LPS treatment. We found that the number of pulmonary Ly6G^high^Ly6C^low^CD11b^+^ PMNs was higher in the αCGRP + LPS-treated group compared to the LPS-only treated group (Extended Data Fig. 9a-9c). Additionally, when iBMDMs were treated with CGRP followed by LPS challenge, we observed that LPS upregulated the expression of chemokines *Cxcl10* and *Cxcl1*, and this effect was enhanced by αCGRP treatment (Extended Data Fig. 9d and 9e). These findings strongly support the idea that PNECs and their product, CGRP, can amplify LPS-induced lung inflammation.

### GABA feedback regulates the production of CGRP in PNECs and vagal sensory neurons

PNECs are main GABA-producing cells in the lung, and GAD1 and ABAT are key enzymes for GABA metabolism ^42^. GAD1 mediates the synthesis of GABA, while ABAT controls its breakdown ^43^. The GABAergic synapse and receptor clustering are regulated by SHISA7, PHF24, LHFPL4, DBI, NRXN1, and NLGN2 ^44^. ACBP (Acyl-CoA Binding Protein, a polypeptide encoded by *Dbi*) binds to GABA receptors and regulates GABA inhibition ^45^. By analyzing our vagal ganglion and lung RNAseq databases, we found that lung *Abat* expression was downregulated under LPS challenge, indicating that GABA is increased in PNECs. Concurrently, *Shisa7*, *Phf24*, *Nrxn1*, *Usp46* were downregulated in PNECs and upregulated in vagal ganglia. *Dbi* expression was shown in the opposite direction (Fig. 7a). These findings indicate that GABA receptors are activated in vagal ganglia and inhibited in PNECs under LPS challenge. Furthermore, we analyzed vagal ganglia GSE192987 ^32^ and PNEC^1^ scRNAseq databases and found that *Gabbr1/2* and *Gabrb3* were expressed by lung-innervated *Trpa1*^+^*Calca^+^* neurons and *Gabrb1/3* and *Gabbr2* were expressed in PNECs^1^ (Extended Data Fig. 10). As such, we hypothesize that PNEC-derived GABA can inhibit the excitability of *Trpa1^+^Calca^+^* neurons and CGRP synthesis in PNECs (Fig. 7b).

**Fig. 7.**
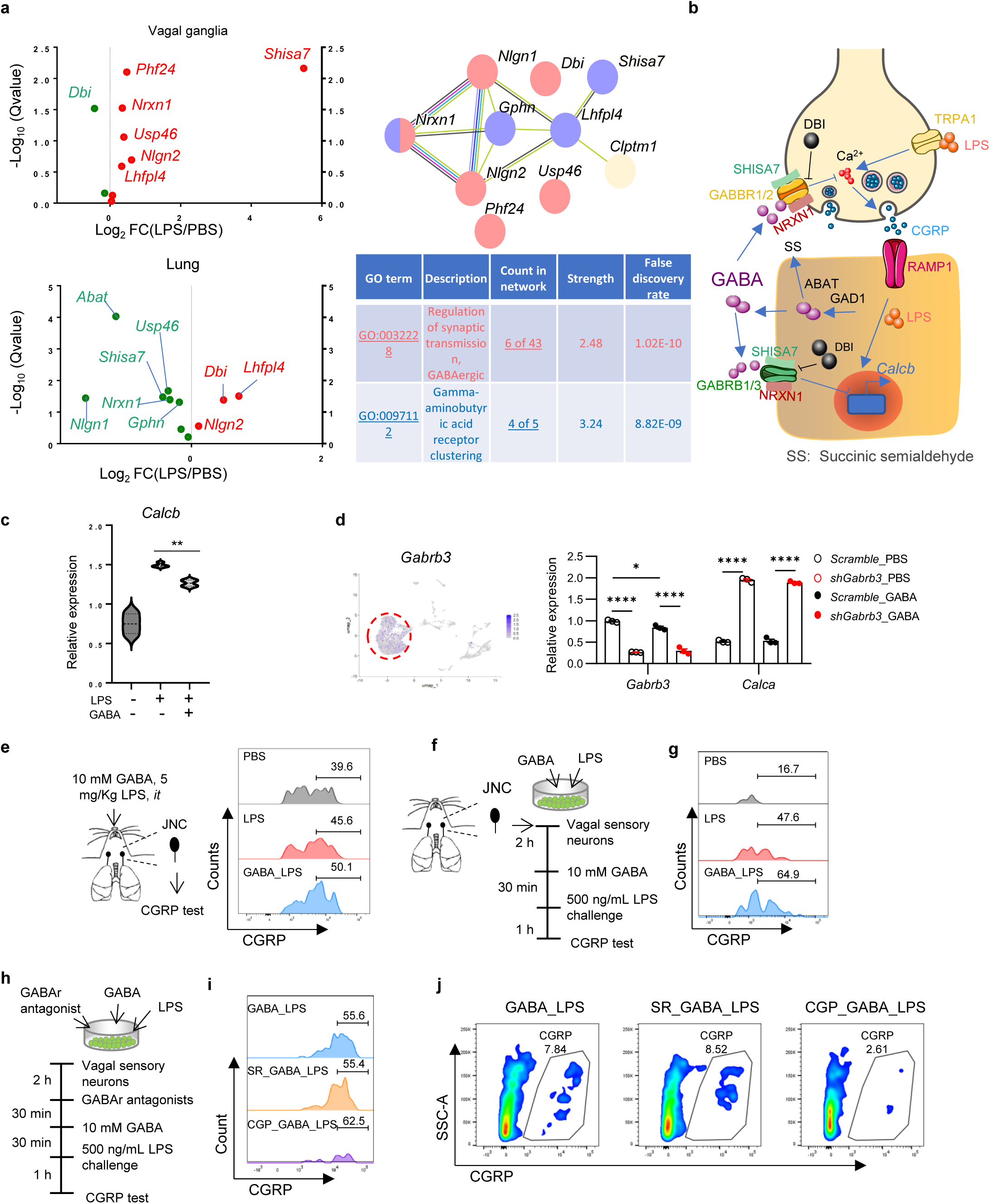
GABA feedback regulates the production of CGRP in PNECs and vagal sensory neurons. (a) Analyze the expression of genes related to GABAergic synapses, GABA metabolism, and its receptors in the RNAseq databases of LPS-stimulated lungs and vagal ganglia. (b) A schematic of the hypothesis that GABA derived from PNECs can inhibit the excitability of *Trpa1^+^Calca^+^* neurons and CGRP synthesis in PNECs. (c) Measurement of *Calcb* gene expression in SCLC cells stimulated with LPS and GABA. (d) Measurement of *Gabrb3* and *Calca* gene expression in Gabrb3-knockdown PNEC cells and control cells after GABA stimulation. (e) The levels of CGRP in the vagal ganglia were measured after GABA and LPS stimulation via endotracheal intubation. (f) Experimental timeline of primary vagal sensory neurons stimulation with GABA and LPS. (g) The levels of CGRP in primary vagal sensory neurons were measured following GABA and LPS stimulation. (h) Experimental timeline of primary vagal sensory neurons stimulation with GABAr antagonist, GABA and LPS. (i and j) The levels of CGRP in primary vagal sensory neurons were measured following treatment with GABA_A_ receptor antagonist SR-95531 (SR) and GABA_B_ receptor antagonist CGP52432 (CGP). One-way ANOVA in (c) and (d). Mean ± SEM. ns, not significant. *p < 0.05, **p < 0.01, and ***p < 0.001.

To test this hypothesis, we pretreated the PNEC cell line with GABA and then stimulated with LPS, and found that LPS increased *Calcb*, and GABA treatment decreased this effect (Fig. 7c). Considering that *Gabrb3* was highly expressed by PNECs^1^, we knocked down this gene and found that *Calca* was upregulated in *Gabrb3*-silenced group (Fig. 7d). These findings support that GABA may inhibit CGRP expression in PNECs. To study whether GABA affects CGRP expression in vagal ganglia, we intratracheally delivered GABA to mouse lungs ^24^ and challenged them with LPS. We found that GABA increased the CGRP expression in vagal sensory neurons from LPS-challenged mice (Fig. 7e). In isolated cell studies, we also found that GABA increased the expression CGRP in LPS-stimulated vagal sensory neurons (Fig. 7f-7g). To compare whether GABA_A_ and GABA_B_ receptor antagonists would prevent the enhanced CGRP expression induced by GABA + LPS, we pretreated neurons with SR-95531 (anti-GABA_A_ receptor) and CGP52432 (anti-GABA_B_ receptor) ^46,47^ (Fig. 7h). We found that blockade of the GABA_A_ receptor did not alter CGRP expression in GABA + LPS-treated neurons compared to the group without antagonist. However, blockage of the GABA_B_ receptor significantly reduced CGRP expression in GABA + LPS-treated neurons (Fig. 7h-7j). These findings suggest that GABA regulates the CGRP release from vagal sensory neurons.

## Discussion

In this study, we identified that αCGRP^+^TRPA1^+^ nodose sensory neurons innervated PNECs and utilized TRPA1 to sense LPS, increasing their αCGRP expression. αCGRP derived from sensory neurons acts on PNECs, not only promoting their CGRP expression but also regulating their proliferation, thereby amplifying pulmonary inflammation responses. GABA (as a major product of PNECs) feedback regulated CGRP production in nodose sensory neurons. Thus, we have uncovered that αCGRP^+^TRPA1^+^ nodose sensory neurons collaborate with PNECs to form neural circuits that regulate LPS-induced lung inflammation (Extended Data Fig. 9f).

It has been demonstrated that LPS can exert fast excitatory actions via TRPA1 in neurons, mediating acute neurogenic inflammation and pain ^16^. Considering the slow activation of traditional immune signaling pathways ^48^, a plausible pattern of recognition is coupling TLR4 receptors with ion channels, such as TLR4-TRPV1, which may explain rapid neuronal activation ^49^. However, antagonism of TLR4 could not completely inhibit LPS-induced neuronal activation, such as calcium influx ^16,50^. In a recent study, bacterial LPS can activate the TLR4 receptor on TRPV1^+^ vagal sensory neurons, transmitting signals to the paraventricular hypothalamic nuclei (PVN), leading to the activation of corticotropin-releasing hormone neurons and inducing sickness behavior ^8^. The vagus nerve response to LPS was assessed by activating the TLR4 receptor, which can increase neuronal sensitivity to capsaicin ^8^. Another in vitro neuronal calcium signaling recording experiment found that TRPA1^+^ neurons in the nodose ganglion can directly respond to LPS stimulation without relying on TLR4^16^. However, in an *in vivo* intestinal neuronal calcium signaling recording experiment, LPS did not directly activate vagal sensory neurons^51^. This issue remains controversial: whether vagal sensory neurons can directly sense LPS in the pulmonary environment in vivo, and which receptor (TLR4 or TRPA1) mediates LPS recognition, is still unclear. In our study, pulmonary LPS-induced vagal excitability was eliminated by deletion of *Trpa1* in the nodose ganglia (Fig. 1k-m), but not by systemic deletion of *Tlr4* (Extended Data Fig. 4a-d and 4i-j). By analyzing scRNAseq datasets of sensory neurons projecting to the lungs ^32^, we found that few *Tlr4*^+^ sensory neurons were presented in QZ2 labeled lung-innervated nodose sensory neurons, and *Trpa1* was 20-fold more expressed than *Tlr4* (Extended Data Fig. 4h). We ligated the middle of the trachea and inserted a catheter to inject LPS into the distal region of the lung. This is done to prevent LPS from regurgitating into the laryngopharynx to activate sensory neurons in the jugular ganglia, where TLR4 is expressed 10-fold higher than in nodular ganglia (Extended Data Fig. 4g). It is estimated that less than 12% of vagal *Tlr4*-positive neurons were reported to be located in the nodose ganglia, and more than 50% of vagal *Tlr4*-positive neurons were located in the jugular ganglia ^52^. *Jin et al.* discovered that LPS cannot directly activate vagal sensory neurons projecting to the gut but instead activates them through inflammatory cytokines ^51^. Indeed, immune cell-derived inflammatory cytokines are important factors in neuronal activation and can also promote the release of CGRP from neurons. However, the time required for immune signaling pathways is much longer than that for neuronal signaling. Moreover, the inflammatory cytokines induced by LPS are complex and diverse, including both pro-inflammatory and anti-inflammatory factors, which activate different populations of sensory neurons. Additionally, we found that, unlike vagal sensory neurons projecting to the gut, those projecting to the lungs exhibit higher expression of TRPA1 and co-express elevated levels of αCGRP (Fig. 2i-2k). These features confer unique functions in pathogen recognition and immune regulation to vagal neurons projecting to the lungs. Thus, we conclude that TRPA1 rather than TLR4 in the nodose sensory neurons senses LPS and couples with MYD88 to increase vagal excitability. This notion is supported by LPS stimulation induces MYD88 aggregation in *Trpa1*-overexpressed neurons (Extended Data Fig. 3b-3d) and deletion of *Myd88* in nodose sensory neurons (*Phox2b^+^*) abolishes LPS-triggered vagal excitability (Fig. 1d-F). Therefore, we focus on the mechanism by which αCGRP^+^TRPA1^+^ nodose sensory neurons directly sense LPS and rapidly mediate neurogenic regulation of lung inflammation.

Different vagal sensory neurons project to various visceral organs^32^. It remains unclear whether vagal sensory neurons, such as those expressing TRPA1 and αCGRP, which project to different organs, have similar inflammatory regulatory functions. A recent study on the vagal-cNST inflammatory circuit revealed that different vagal sensory neurons can respond to different inflammatory signals. TRPA1^+^ neurons respond to anti-inflammatory factor IL-10, while CGRP^+^ neurons respond to pro-inflammatory factor IL-6 and IL-1*β*, thereby suppressing systemic inflammatory responses^7^. This anti-inflammatory effect originates from the activation of sensory neurons, whereby signals are transmitted to DBH^+^ neurons in the cNST, which mediate inflammatory regulatory reflexes, such as through the cholinergic anti-inflammatory pathway^9^. In our study, we propose that TRPA1 expressed in αCGRP^+^ sensory neurons can directly sense LPS and make neurons synthesize αCGRP to influence PNECs. Through analyzing the database by Chang and colleagues^32^, we found that vagal sensory neurons projected to the lungs are very different from those projecting to abdominal organs (Fig. 2j). Compared to esophagus, stomach, duodenum, colon, and pancreas, QZ2 labeled lung-innervated nodose sensory neurons have the highest number of *Trpa1^+^Calca^+^* cells (Fig. 2i). Of these organs, there are very few single *Calca^+^*sensory neurons innervating the lungs (Extended Data Fig. 11a and 11b). Deletion of *Trpa1* or *Calca* in sensory neurons had the same effect on CGRP expression in PNECs and lung inflammatory responses, suggesting that *Trpa1^+^Calca^+^*sensory neurons, rather than single *Trpa1^+^* or *Calca^+^*sensory neurons, can mitigate their effects on PNECs (Fig. 5).

Vagal nodose sensory neurons project various types of sensory terminals at the NEBs, among which *Pvalb*^+^ neurons are responsible for responding to airway closure ^1^, but the functions of nodose TRPA1^+^αCGRP^+^ neurons remain largely unresolved. We observed that in larger airway, αCGRP^+^ neurons of the NG project to PNECs. The nerve endings of these neurons are primarily located beneath the airway epithelium, especially projecting below the NEB structure, rather than extending internally into the NEBs, suggesting that these αCGRP^+^ nodose sensory neurons may not contact with LPS. However, in the distant airway, CGRP^+^ nerve fibers can extend into the alveolar space, indicating that nerve endings in these areas may sense LPS (Fig. 3).

One important finding in this study is that nodose *TRPA1^+^αCGRP^+^* neurons innervate PNECs and release αCGRP to promote CGRP production in PNECs in response to LPS challenge. PNECs are one of the epithelial cell types lining large and small airways that form a tiny cellular population (0.4% of total airway epithelial cells or 0.01% of all lung cells)^53^. NEBs frequently populate diametrically opposed positions to the bifurcation points of branching airways^53^. For defense against bacterial invasion, *Trpa1^+^Calca^+^* nodose sensory neurons can sense LPS to produce CGRP and amplify its effects via PNECs. We used EpCAM^+^NCAM1^+^CD45^−^ cells to define PNECs, and 80∼90% of cells are CGRP^+^ in this cell population. In our study, deletion of *Trpa1* or *Calca* in sensory neurons reduced CGRP expression in PNECs from LPS-challenge mice. Our study supports the activation of TRPA1^+^αCGRP^+^ nodose sensory neurons as a mechanism to amplify LPS-induced lung inflammation. However, several reports suggest that activation of nociceptor sensory neurons can suppress lung inflammation^54–56^. Nociceptors are a diverse group of sensory neurons that include various subtypes capable of releasing CGRP, SP, VIP, and other neuronal mediators^48,57^. Notably, CGRP^+^ sensory neurons are predominantly localized in the jugular ganglion^57^, while CGRP+ neurons in the nodose ganglion are often overlooked. CGRP can be further divided into αCGRP (encoded by *Calca*) and βCGRP (encoded by *Calcb*), but previous studies have often referred to them collectively as “CGRP.” This distinction is crucial, as the expression patterns of these genes differ significantly in vagal sensory neurons (Extended Data Fig. 4e and 4f). For instance, βCGRP^+^ vagal sensory neurons have been shown to suppress group 2 innate lymphoid cell function and allergic airway inflammation^56^. However, the role of nodose CGRP^+^ sensory neurons, particularly αCGRP, in LPS-induced pneumonia remains unclear. Most of these studies utilized *Trpv1*^Cre^ and *Nav1.8*^Cre^ mice to investigate sensory neuron functions. TRPV1 and Nav1.8 are markers of nociceptive neurons, encompassing a broad range of neuron types, not limited to αCGRP-positive neurons. Additionally, our study focuses on αCGRP. Furthermore, using *Nav1.8^Cre^/Rosa26^DTA^* mice results in the absence of systemic nociceptive neurons from birth, or systemic depletion of nociceptive neurons via intraperitoneal injection of diphtheria toxin (DTA). In our study, we used *Calca^CreERT2^/Rosa26^DTR^* mice and achieved specific ablation of αCGRP^+^ neurons within the nodose ganglion through localized injection of DTA directly into the nodose ganglion. Due to differences in the target cells, sensory neurons may exert different regulatory effects. Our focus is on αCGRP^+^ neurons targeting PNECs. PNECs themselves contain various bioactive substances that, together with CGRP, collectively regulate lung inflammatory responses.

At the same time, compared to PNEC cell lines treated with αCGRP alone, both protein and transcriptional levels of CGRP were increased in the LPS + αCGRP treatment group, suggesting that LPS-induced signaling is essential for the αCGRP-enhanced CGRP expression in PNECs. Unlike previous reports^38–40^, PKA signaling may negatively regulate CGRP expression in PNECs, as evidenced by higher CGRP levels in cell lysates and supernatants in the H89 (PKA inhibitor) + αCGRP treatment group than in the αCGRP treatment group (Extended Data Fig. 8). Whether PKA signaling is involved in LPS stimulation-enhanced CGRP expression is worthy of further investigation. As previously reported, the release of CGRP from respiratory vagal afferents is critical to bacterial clearance in lungs ^55^.

Taken together, we have identified a population of *Trpa1*^+^*Calca*^+^ sensory neurons that project to the lungs, detect intrapulmonary LPS, trigger vagal electrical excitations, and interact with PNECs through CGRP. Meanwhile, PNECs also release GABA, providing negative feedback inhibition on vagal CGRP. Therefore, we have established a nodose ganglion-pulmonary neuroepithelial circuit that senses and regulates lung inflammatory responses.

## Acknowledgements

We thank all members of the Su lab for their helpful comments and discussion. We thank Prof. Pao-Tien Chuang (University of California San Francisco) for *Calca^CreER^* mice. This work was supported by NSFC programs (82241042, 82495200/02, 82400103); National Key Research and Development Program of China (No.2022YFC2304702); Science and Technology Commission of Shanghai Municipality (20DZ2261200); China Postdoctoral Science Foundation (2024M750522).

## Author contributions

J.C. and X.S. conducted planning and conceptualization; X.S. provided supervision; J.C. and S.X. analyzed data and organized it into Fig.; J.C. and C.Z. performed sequencing; S.X. analyzed vagus nerve activity; J.C., S.X., and R.T. conducted animal experiments; J.C. conducted cell experiments; J.C., S.X., and Z.L. conducted confocal imaging; J.C. and S.X. conducted flow cytometry and ELISAs; J.C. and Z.L. prepared the viral vector; J.C., S.X., Z.L., R.T., C.Z., X.C., Y.Z., Y.M., and R.W. developed the methodology; key resources were provided by H.S., S.W., X.Z., P.S., H.J., and Y.S.; funding acquisition was managed by X.S. and J.C; J.C. wrote the original draft; and X.S. reviewed and edited the manuscript.

## Competing interests

The authors declare no competing interests.

## Methods

### Mice

C57BL/6J mice were purchased from Charles River Laboratories. *Tlr4* knockout (*Tlr4^−/−^*) mice and *Rosa26-stop(flox)-DTR*, Ai9 strain *Rosa26-stop(flox)-tdTomato* mice were also obtained from Jackson Laboratory. *Phox2b^Cre^* mice were provided by S. Wang (Hebei Medical University). *MyD88^flox^* mice were provided by X. Zhang (Shanghai Institute of Immunology, CAS). *Ascl1^CreERT2^* mice were provided by P. Sui (Shanghai Institute of Biochemistry and Cell Biology, CAS). *Calca^CreERT2^* mice were provided by H. Song (Zhejiang University).

For labeling of nodose ganglion sensory neurons, *Phox2b^Cre/Cre^* mice were crossed with *Rosa26^stop(flox)−tdTomato^* mice to generate nodose ganglion neuron lineage-labeled mice (*Phox2b^tdTomato^*). For nodose ganglion-specific MyD88 deletion experiments, *Phox2b^Cre/-^*; *MyD88^fl/fl^* mice were crossed with *MyD88^fl/fl^*mice to generate mice with nodose ganglion-specific MyD88 deletion (*MyD88^Phox2b^)* or control (*MyD88^fl/fl^*) littermates. For Tlr4 knockout mouse experiments, Tlr4^+/−^ heterozygous mice were paired to produce wild-type *Tlr4^+/+^*(*Ctrl* mice) and Tlr4 knockout *Tlr4^−/−^* (*Tlr4^KO^*) mice. For Calca lineage neuron depletion experiments, *Calca^CreERT2/-^*; *Rosa26^stop(flox)−DTR/stop(flox)−DTR^* mice were crossed with *Rosa26^stop(flox)− DTR/ stop(flox)−DTR^*mice to generate Calca lineage neuron-depleted mice (*NG^CalcaABLATE^*) and control *CalcaCreERT2-/-; Rosa26^stop(flox)−DTR/stop(flox)−DTR^* (*Ctrl*) littermates. Then, DT was injected into the nodose ganglion to achieve the depletion of Calca lineage neurons. For vagal ganglion-specific TRPA1 deletion experiments, we used C57BL/6J mice and injected *AAV-sgTrpa1* (*NG^Trpa1KO^* mice) or *AAV-Ctrl* (*Ctrl* mice) into the nodose ganglion. For the experiment of PNEC ablation, *Ascl1Cre^ERT2/-^; Rosa26^stop(flox)−DTR/stop(flox)− DTR^* mice were crossed with *Rosa26^stop(flox)−DTR/stop(flox)−DTR^* mice to generate Ascl1 lineage-ablated mice (*Ascl1^CreERT2/-^; Rosa26^stop(flox)−DTR/stop(flox)−DTR^* mice or *Ascl1^ABLATE^* mice) and control littermates (*Ascl1^CreERT2^-/-; Rosa26^stop(flox)−DTR/stop(flox)−DTR^* or *Ctrl mice*).

All animal experiments complied with the animal care standards were approved by the Committees on Animal Research of the Shanghai Institute of Immunity and infection, Chinese Academy of Sciences, China. These mice were bred and maintained in an SPF animal facility at Shanghai Institute of Immunity and infection. The mice were housed with free access to food (ad libitum) and water in 12-h dark/light cycle. Both male and female mice were used in all experiments. Anesthesia was induced with an intraperitoneal (ip) injection of Avertin (1.25%). The number of mice used is as indicated in Fig. legends; 3 or more mice were used per experimental condition.

### Mouse animal models

The mice were anesthetized by inhalation of isoflurane. They were intratracheally injected with either the indicated concentration (5 mg/kg) of LPS dissolved in PBS or an equal volume of PBS. Similarly, CGRP (6.5 μg) or PBS was intratracheally delivered.

### Cells

Cell cultures were maintained in a humidified atmosphere at 37°C with 5% CO2. SCLC cells were cultured as previously described ^35^. Briefly, cells were cultured in RPMI 1640 medium supplemented with 10% FBS (Gibco) and 1% penicillin/streptomycin (Hyclone). After observing the cell status and confluency, cells were harvested by centrifugation at 800 rpm for 5 minutes following 1-2 days of culture. The supernatant was discarded, and the cell pellet was resuspended in 2 ml of fresh medium. Cells were then gently pipetted multiple times to dissociate cell aggregates and obtain a single-cell suspension, which was used for subsequent experiments.

Neuro-2a (ATCC CCL-131) cells were cultured in EMEM medium supplemented with 10% FBS, and 1% penicillin/streptomycin. Trypsin-EDTA solution (Gibco) was added to obtain a cell suspension for subsequent experiments.

### Viruses

rAAV vectors were generated in 293T cells by transfecting packaging plasmids (pHelper, pRep/Cap) and the ITR-containing vector plasmid in a 1:1:1 molar ratio. The supernatant was collected at 72 and 120 hours after transfection, and the cells were collected at 120 hours after transfection. Viral particles were concentrated by iodixanol density gradient centrifugation at 288,000 × g for 3 hours at 10°C. AAV titers were quantified by qPCR ^58^. Primer sequences for qPCR are listed in the table. For lentivirus production, HEK293 cells in a 10 cm dish were transfected with 4.5 µg of the psPAX2 plasmid (Addgene#12260), 1.5 µg of the pMD2.G plasmid (Addgene#12259), and 6 µg of the vector plasmid. Lentiviruses were harvested from the supernatant at 48 and 72 hours after transfection.

### Plasmids construction

To label MyD88 in live cells ^59^, mouse MyD88 was modified with EGFP at the C-terminus using overlap PCR. The MyD88-GFP fragment was then cloned into the lentiviral backbone pWPI-IRES-puro. To overexpress Trpa1, mouse Trpa1 was fused with IRES-mCherry and inserted into a plasmid driven by the CMV promoter (CMV-Trpa1-IRES-mCherry). The AAV plasmids for CRISPR guidance were modified from pX601-saCas9 (Addgene#61591). In brief, the saCas9 fragment was fused with the hsyn promoter, WPRE, and bGH (poly A) to generate the hsyn-saCas9-WPRE-bGH fragment. This fragment was then inserted between the two ITRs of the pX601 donor plasmid to generate pXN-hsyn-saCas9. For the sgRNA vector, saCas9 in pX601 was replaced with mCherry. sgRNA oligos were designed using CRISPick (https://portals.broadinstitute.org/gppx/crispick/public) and cloned into the U6-sgRNA scaffold. The U6-sgRNA units were then amplified by PCR and cloned into the pX601-mCherry backbone using Gibson assembly, as described previously ^60^.

### Tamoxifen administration

For Calca lineage neuron depletion experiments, *Calca^ABLATE^* mice and *Ctrl* littermates were anesthetized with 1.5-2.0% isoflurane in air. The vagal ganglion was exposed following a midline incision in the neck. Then, 20 ng of diphtheria toxin (DT) in 120 nL of PBS was injected into the NG using a nanoinjector (Drummond Scientific Company). After a 2-week recovery period, the mice underwent further experiments.

For the experiment of PNEC ablation, *Ascl1^ABLATE^* mice and *Ctrl* mice were intraperitoneally injected with 100 μL of tamoxifen (Sigma) (20 mg/mL in corn oil) for 5 consecutive days. Experiments were performed 7 days after the last administration.

### Bronchoalveolar lavage in Mice

Collect bronchoalveolar lavage fluid (BALF) as previously described ^61^.Mice were euthanized under avertin anesthesia. The neck skin was cut, and blunt dissection was performed with forceps to expose the trachea. A small incision was made on the trachea, and care was taken not to cut it completely. A 1 mL syringe was used to aspirate 1 mL of PBS, and then replaced with a blunt needle. The needle was inserted through the small incision into the trachea. A surgical knot was made with a surgical thread to prevent liquid leakage. The lavage was repeated three times. Cells in the lung lavage fluid were obtained by centrifugation, and the supernatant was collected to measure protein and inflammatory cytokine concentrations.

### Cytokine measurement

The collected procedures for Bronchoalveolar lavage (BALF) were described above. TNF-α concentrations were analyzed in the BALF using the TNF-α ELISA Kit following the manufacturer’s instructions.

### Isolation of mouse lung primary cells

The mice were euthanized, and the lungs were dissected and dissociated as previously described ^62^. After euthanasia, the mice were fixed on a surgical board. The chest cavity of the mice was opened, and the heart was exposed. The left atrium was excised, and the heart was perfused. A 10 mL disposable syringe was inserted into the right ventricle from the apex of the heart, and 10 mL of chilled (4°C) PBS solution was slowly and evenly injected to remove the blood from the body. Subsequently, 1 mL of tissue digestion solution (1 mg/mL collagenase (sigma) /2.2 mg/mL dispase mixed solution (Roche)) was injected into the lungs through the trachea. After tying off the trachea, the lung tissue was removed and placed in a 15 mL centrifuge tube containing 1 mL of tissue digestion solution, and digested in a shaking water bath at 37°C for 30 minutes. After digestion, the lungs were carefully aspirated using a Pasteur pipette to obtain a single-cell suspension of whole lung cells. The lung cells were washed, subjected to red blood cell lysis (Biolegend), and filtered through a 70 μm nylon mesh (BD). The lung cells were resuspended in RPMI 1640 medium (10% FBS and 1% penicillin/streptomycin), counted, plated in cell culture plates, and incubated for 24 hours before further experiments.

### Antibodies

The primary antibodies and the secondary antibodies used in the procedure include:Rabbit anti-mouse TRPA1 (Proteintech, 19124-1-AP), Rabbit anti-mouse CGRP-I+CGRP-II (Abcam, b283568), Rat anti-mouse NCAM1 BV421 (BD, 78094), Rat anti-mouse NCAM1 BV605 (BD, 809220), Rat anti-mouse CD11b PEcy7 (Biolegend, 101216), Rat anti-mouse F4/80 BV421 (BD, 565411), Rat anti-mouse Ly6G AF488 (Biolegend, 127626), Rat anti-mouse EpCAM PE (Biolegend, 118205), Rat anti-mouse EpCAM APC (Biolegend, 118213), Rat anti-mouse CD45 FITC (Biolegend, 123107), Rat anti-mouse CD45 APCcy7 (Biolegend, 103116), Rat anti-mouse Ly6C APC (Biolegend, 128016), Rat anti-mouse Ly6C BV421 (Biolegend, 128031), Rat anti-mouse TNF-α PE (Biolegend, 506306), Rat anti-mouse NCAM1 APC (Abcam, ab237383), Donkey anti-rabbit IgG AF647 (Biolegend, 406414), Goat anti-rabbit IgG AF700 (Invitrogen, A-21038), Goat anti-Rabbit CoraLite488 (Proteintech, SA00013-2), Goat anti-rabbit Cy3 (Servicebio, GB21303), Rabbit anti-GAPDH (CST, 5174), Rabbit anti-β-Actin (CST, 4967), Goat anti-rabbit IgG HRP (Jackson Immuno, 111-035-144).

### Flow cytometry

The detailed procedures were described previously^61^. The isolation of lung cells for flow cytometry followed the previously described steps. For flow cytometry staining of lung cells, the following sequential steps were performed: viability staining, surface staining, and intracellular staining. 0.1 µl of Zombie Yellow or Zombie NIR (Biolegend) was added to each sample to assess cell viability. The samples were then incubated at room temperature for 30 minutes, followed by two washes with FACS buffer and centrifugation at 1500 rpm. After preincubating for 15 minutes with anti-mouse CD16/32 antibodies (Biolegend), single-cell suspensions were stained for 30 minutes on ice with surface antibodies at a 1:100 dilution in FACS buffer, then washed twice. For intracellular staining of cytokine TNF-α+(Biolegend) and CGRP+ cells, cells were fixed and permeabilized with the Fixation/Permeabilization Kit (BD) for 20 minutes at 500g, followed by incubation with intracellular antibodies at a 1:100 dilution in Permeabilization Buffer for 30 minutes, then washed twice. For CGRP staining, indirect labeling was performed using the secondary antibody Goat anti-rabbit AF700. The cells were washed twice and resuspended in FACS buffer for analysis by flow cytometry. Debris and aggregates were excluded, and live cells were analyzed using an LSRFortessa flow cytometer (BD Biosciences). Data were analyzed using FlowJo software (Tree Star Inc).

The isolation of neurons from the vagal ganglia followed the same procedures as described earlier for sensory neuron isolation and culture. The staining process was similar to that of lung cells, including viability, surface, and intracellular staining, with a modification of the centrifugation speed to 800g.

### RNA isolation, reverse transcription, and RT-PCR

Total RNA was isolated from cells or homogenized lungs and DRG using TRIzol reagent (Invitrogen) according to the manufacturer’s instructions. RNA was quantified and reverse-transcribed to generate cDNA using a reverse transcriptase kit (Yeasen), which was followed by quantitative real-time polymerase chain reaction analysis. RT-PCR was performed using the QuantStudio™ 6 Flex system (ABI) with SYBRGreen Mix (Yeasen). The fluorescence values were measured by the instrument to determine the cycle threshold (Ct). The 2−ΔΔCT method was used to analyze the target gene expression, and the Gapdh mRNA levels were used as an internal control for normalization in each sample. The primers used in the procedure include: *Gapdh* 5’-CCCACTAACATCAAATGGGG-3’(forward) and 5’- CCTTCCACAATGCCAAAGTT-3 ′ (reverse); *Cxcl1* 5 ‘ -ACTGCACCCAAACCGAAGTC-3 ‘ (forward) and 5 ‘ - TGGGGACACCTTTTAGCATCTT -3′(reverse); *Ccl2* 5’- TTAAAAACCTGGATCGGAACCAA -3’(forward) and 5’- GCATTAGCTTCAGATTTACGGGT -3 ′ (reverse); *Il6* 5 ‘ -CCCCAATTTCCAATGCTCTCC-3 ‘ (forward) and 5 ‘ - CGCACTAGGTTTGCCGAGTA-3′(reverse); *Calca* 5’-CCTTTCCTGGTTGTCAGCATCTTG -3’(forward) and 5’ - CTGGGCTGCTTTCCAAGATTGAC-3′(reverse); *Calcb* 5’-CTCTCAGCACGATATGGGTCC-3’(forward) and 5’ - GCAAGAGATGTTTTTCCTGGTCG-3′(reverse);

### Immunofluorescence

Animals were euthanized by intravenous injection of Avertin (1.25%). The chest cavity of the animals was then opened for standard transcardial perfusion with PBS buffer for 5-10 minutes at room temperature until the blood was washed out, followed by perfusion with 4% paraformaldehyde (PFA) for 10-20 minutes (Servicebio). The DRG, nodose ganglia, and lung were dissected and post-fixed overnight at 4°C. The ganglia were fixed for 1 hour at 4°C. These tissues were then incubated overnight at 4°C in 30% sucrose/PBS (Sigma), embedded in Tissue-Tek OCT compound (Sakura), and frozen using liquid nitrogen. Tissues were sectioned into 10 μm to 30 μm slices using a cryostat. The slices were blocked with blocking solution (5% BSA, 0.1% Triton X-100 in PBS) at room temperature for 1 hour, followed by overnight incubation with primary antibodies at 4°C. After washing, the sections were incubated with secondary antibodies for 1 hour. Following washing with PBS, the slides were mounted using DAPI mounting medium (Abcam).

For cell experiments, cells were plated on coverslips. After stimulation, the culture medium was discarded, and cells were washed twice with PBS. Cells were fixed with 500 μl of 4% PFA (Servicebio) for 20 minutes. After two washes with PBS, cells were permeabilized with 0.1% Triton X-100 for 15 minutes at room temperature. Blocking was performed with blocking solution (5% FBS, 0.1% Triton X-100 in PBS) for 20 minutes. Cells were then incubated with primary antibodies overnight in a humidified chamber. After washing, cells were incubated with secondary antibodies at room temperature for 60 minutes. Following three washes with PBS, cell nuclei were stained with DAPI and coverslips were mounted (Abcam).

### Detecting Myd88 oligomerization

We first transfected Neuro-2a cells with *Myd88-GFP* to establish the *Neuro-2a-MyD88-GFP* cell line. Next, we transfected the *Neuro-2a-MyD88-GFP* cells with *Trpa1-IRES-mCherry* or a control plasmid, *Cas9-IRES-mCherry*. After 30 minutes of LPS stimulation, we collected samples from Neuro-2a cells overexpressing Trpa1 (*Trpa1^ov^*^er^) and control cells (*Ctrl*) and acquired data using fluorescence microscopy. We then used HALO software (Indica labs) to automatically count the number of MyD88-GFP puncta larger than 0.5 μm² per individual cell.

### Ganglia clearing by CUBIC

Advanced CUBIC protocols for tissue clearing were introduced as described previously ^63^. Briefly, mice were anesthetized and transcardially perfused with cold PBS to remove the blood from the organs as much as possible. Subsequently, they were perfused with 4% PFA. The ganglia were dissected and fixed in 4% PFA, with shaking at 4°C for 1 hour. After washing with PBS three times, the samples were immersed in Reagent-1 diluted with 1/2 water (25 wt% urea, 25 wt% Quadrol, 15 wt% Triton X-100) and shaken at 37°C for 2 hours. The 1/2-diluted Reagent-1 was discarded. Reagent-1 was replaced, and primary antibodies were added, then gently shaken at 37°C for 1 day. After that, the samples were incubated overnight with secondary antibodies in Reagent-1. The samples were degassed using a vacuum desiccator. Then, the samples were immersed in Reagent-2 diluted with approximately 1/2 PBS (25 wt% urea, 50 wt% sucrose, 10 wt% triethanolamine), shaken at 37°C for 2 hours, and then treated in Reagent-2 for one day. Clear ganglia were imaged using a confocal microscope (Olympus).

### Vagus nerve recording

The vagus nerve recordings were performed as described previously ^64^. Briefly, mice were induced under anesthesia with 3% isoflurane in 100% air at a flow rate of 0.9 L/min for 3 min. Subsequently, the mice were maintained under anesthesia with 1.5% isoflurane and placed on a heating pad to maintain a body temperature of around 37°C. To isolate the cervical vagus nerve, the neck area was cleaned with iodine, and a midline cervical incision was made from the level of the larynx to the sternum. Submandibular salivary glands and the muscles on the trachea were separated to expose the trachea. A small dosing tube was inserted into the lung by perforating the trachea with a 29G needle and placing the dosing tube approximately 7 mm deep into the trachea. The right cervical vagus nerve was carefully separated from the artery and desheathed by gently removing the thin connective tissue surrounding the nerve under magnification. The vagus nerve was then placed on two hook electrodes with mineral oil to insulate the nerve and the surrounding tissue. A ground electrode was inserted into the mouse’s tail. The PowerLab 4/26 data-acquisition system (PowerLab; ADINSTRUMENTS, Inc.) was used to record signals from the vagus nerve. Recordings were sampled at 40 kHz, filtered at 120 Hz, and amplified 100 times to ensure the signals were stable enough for long-term recording. Following the acquisition of baseline activity (10 min), 10 μL of LPS (10 mg/mL) or PBS (as control) was injected into the lung via the dosing tube, and recordings continued for 20 min post-injection. The analysis process was previously described^64^. In brief, the signal in mat format was exported from LabChart (LabChart; ADINSTRUMENTS, Inc.) and analyzed in MATLAB. The signals were decomposed into individual spikes, and the action potentials were detected using a threshold determined by a smallest of constant false-alarm rate (SO CFAR) filter. The t-SNE method was used to perform dimensionality reduction of all spike data, and the DBSCAN method was used to perform cluster analysis of all spikes. Then, all spikes were clustered into different groups based on their waveforms. All significant waveforms were counted, and their distribution over time was analyzed after normalization.

### Calcium imaging and analysis of the jugular-nodose ganglia

*AAV9-CMV-GCaMP6s* was injected into the jugular-nodose ganglion of a 4-week-old mouse to express GCaMP6s in the neurons of the ganglion. After a 4-week incubation period, the animals underwent a calcium signal recording procedure. Mice were administered isoflurane anesthesia via a mask with an airflow rate of 0.8 L/min at a 2% isoflurane concentration. They were placed on a heating pad to maintain body temperature. To isolate the nodose ganglia, the neck area was disinfected with iodine, and a midline cervical incision was made from the level of the larynx to the sternum. Initially, the vagal nerve was isolated, and a dosing tube was inserted into the trachea, as previously described. Subsequently, two blunt hooks were used to detach the muscles and expose the jugular-nodose ganglia. A pair of micro dissecting spring scissors was then used to excise the nodose on the rostral side. Once separated, PBS was used to keep the nerve and ganglia moist. The isolated ganglia were placed on a platform made from a coverslip, held in place by a magnetic stand. The miniscope was also held by a magnetic stand with a lifting rod to adjust the distance between the lens and the ganglia^65^. The miniscope was adjusted to capture 20 images per second (20 fps) and recording was initiated. After approximately 3 minutes of baseline ganglia recording, 25 μL of LPS (10 mg/mL) or PBS was injected into the lung through the dosing tube. Recording was continued for about 5-7 minutes post-stimulation. All raw video data were analyzed using the open-source analysis pipeline named Calcium Imaging Analysis (CaImAn) in Matlab^66^. The fluorescence intensity of the region of interest in the videos was extracted using ImageJ. The first 3 minutes of the videos were considered as the baseline for GCaMP6s. Fold change calculations based on the baseline fluorescence intensity were performed, and the data was ultimately plotted in the form of a heatmap.

### Genetic ablation of αCGRP^+^ neurons in NG

5-week-old *DTR^CalcaCreERT2^* mice or *DTR*^fl/fl^ mice (as controls) were intraperitoneally injected with 100 μL of tamoxifen (Sigma) (20 mg/mL in corn oil) for 5 consecutive days. After two weeks, 20 ng DT in 120 nL PBS was injected into nodose ganglion with nanoinjector (Drummond Scientific Company). Mice were anesthetized with 1.5-2.0 % isofluorane with air (RWD). The vagal ganglion was exposed after a midline incision in the neck. Waiting for 2 weeks recovery, the mice were performed experiments.

### AAV microinjection and AAV-guided Trpa1 knockout in JNC

To knock out Trpa1 in JNC neurons, we generated pX601-mCherry(sgTrpa1), which contains two U6-sgRNA units targeting Trpa1 (sgTrpa1-1: GCTTAACGATGTCAGTGGCTCC; sgTrpa1-2: GTATACCGGAAGTAGTGATATT). The control plasmid contains two empty U6-sgRNA units (sg-empty: GGAGACCACGGCAGGTCTCAG). These vectors were packaged into AAV-PHP.S serotype with hsyn-driven saCas9 (pXN-hsyn-saCas9) to generate AAV-Cas9 and AAV-sgRNA. The AAV was diluted to 10^12^ vg/ml after titration. Five-week-old mice were anesthetized with isoflurane and their limbs were immobilized. A 1 cm incision was made in the ventral neck area skin. Under a dissecting microscope, the fascia and adipose tissue were bluntly dissected, taking care to avoid damaging surrounding blood vessels, to expose the vagal ganglia clearly. Using a Nanoject III Injector (Drummond), 120 nl of AAV-Cas9 and 120 nl of AAV-sgRNA was slowly injected into the vagal ganglia. The injector needle was left in place for an additional 5 minutes before slow withdrawal, allowing viral particles to diffuse and be absorbed at the injection site.

### Short hairpin RNA (shRNA)–mediated RNA interference

Short hairpin RNA (shRNA)-mediated RNA interference was employed, with shRNA sequences designed based on the target gene and synthesized as corresponding stem-loop primers (Sangon Biotech). These primers were annealed and inserted into the pLKO.1 plasmid between the EcoRI and NheI restriction sites (CST). The shRNA pLKO.1 construct was introduced into competent E. coli DH5α (Tsingke) for plasmid amplification and extraction. The resulting plasmid was then co-transfected with VSV-G and Δ 8.9 plasmids into 293T cells. After culturing, the produced lentivirus was used to infect PNEC cells, and the infected cells were selected with puromycin to establish stable cell lines. Th knockdown assay primer sequences were, Scramble CAACAAGATGAAGAGCACCAA; Prkaca CGAGTAACTTTGACGACTATG; Prkacb GTCATGTTGGTGAAGCATAAA.

### CGRP release of SCLC cultures and plasma

SCLC cells were cultured in a 12-well plate (2×10^5^ cells per well) following the above-mentioned method for one day. The cells were stimulated with 100nM CGRP (Phoenix pep) for 30 minutes. After stimulation, the media was aspirated and the cells were washed three times with fresh media. The CGRP concentration in 100 μL of culture supernatant was measured using a CGRP EIA Kit (Cayman Chemicals). For CGRP release in plasma, blood was collected from mice and centrifuged at 500xg for 10 minutes. 100 μL of plasma was collected, and the CGRP concentration was measured using a CGRP EIA Kit (Cayman Chemicals).

### Western blot

The samples were collected by scraping as described above. After quantification by the BCA protein assay (Thermo Scientific), equal amounts of protein were mixed with loading buffer and boiled at 100°C in a water bath for 10 minutes. The prepared protein samples were loaded onto SDS-PAGE gels for electrophoresis. Initially, electrophoresis was performed at 80 V for 30 minutes for gel concentration, followed by adjusting the voltage to 120 V until the ideal separation was achieved. For protein transfer, a wet transfer method was employed to transfer the proteins from the gel to a PVDF membrane (Millipore). The transfer was conducted at 200 mA for 60 minutes. After transfer, the membrane was blocked with 5% skim milk (BD) at room temperature for 1 hour, followed by three washes with TBST. Subsequently, the membrane was incubated with the primary antibody overnight at 4°C. After the primary antibody incubation, the membrane was incubated with the secondary antibody, which was an HRP-conjugated antibody (Jackson Immuno), at room temperature for 1 hour. Following the secondary antibody incubation, the membrane was washed three times with TBST, each for 10 minutes, and then subjected to chromogenic detection using an ECL substrate (Yeasen). Data were collected using a Tanon chemiluminescence imaging system. The bands were analysed and quantitated using Image J software.

### Single-cell isolation of vagal sensory neurons

Vagal sensory neurons were isolated as previously described ^67^. Briefly, Ganglion Dissociation Solution (GaDS) was prepared by combining Advanced DMEM/F12, Glutamine, HEPES, N2 supplement, B27, and NGF. Mice were anesthetized with avertin via intraperitoneal injection. Following euthanasia, the vagal ganglia were dissected out and two local cuts were made to facilitate dissociation. The ganglia were transferred into a centrifuge tube (ganglia from 3 mice were combined into one sample to ensure flow cytometric analysis). The ganglia were washed three times with PBS, allowed to settle at the bottom of the tube, and then the PBS was removed. Next, 1 mL of GaDS and 22 μL of digestion enzymes (collagenase I/II and protease mixture (Roche), 2.5 mg/mL, in H_2_O) were added to the tube. The tube was then incubated at 37°C for 30 minutes with gentle shaking every 15 minutes and tapping it lightly. After digestion, the GaDS solution was removed and the ganglia were washed twice with 1 mL of DPBS. Subsequently, 200 μL of GaDS was added to the tube, and the solution was pipetted up and down until no intact tissue pieces were visible, while avoiding bubble formation. The dissociated cells were passed through a 70 μm cell strainer. Carefully, 300 μL of cells were layered onto the percoll density gradient and centrifuged at 2,900×g for 10 minutes. After centrifugation, the top 700 μL of solution containing a large amount of cell debris was removed and discarded. Then, 700 μL of fresh GaDS was added to the remaining cell-containing solution, and the solution was pipetted up and down several times to mix. After centrifuging at 2,900×g for 15 minutes, the supernatant was carefully removed and discarded, leaving the cell pellet. Finally, the cells were resuspended in 300 μL of GaDS, and the sample was kept on ice for further experiments.

### Retrograde Tracing of the vagal subgroups projecting to the lung

Fast Blue Retrograde Tracing was performed as previously described ^67^. Briefly, anesthetize mice with intraperitoneal injection of avertin. Place the anesthetized mouse on a 45° inclined platform with a heating pad and suspend the mouse by its incisors on a fixed line on the platform to maintain a supine position. Position a cold light source above the trachea. Gently grasp the mouse’s tongue with flat-tipped forceps and slightly move the tongue to expose the translucent opening of the trachea. Insert a 22 G blood collection needle into the trachea to a depth of 0.5-1 cm, withdraw the needle, and slowly drip 40 μL of Fast Blue solution (0.4 mM Fast Blue, 1% DMSO, in PBS) into the blood collection needle. With each breath, the mouse will slowly inhale the Fast Blue solution into its lungs. To prevent the dripping material from overflowing from the trachea, keep the blood collection needle in place for 10 seconds before withdrawing it, and continue to suspend the mouse on the inclined surface for at least 1 minute. If respiratory arrest occurs, chest massage can be performed for cardiopulmonary resuscitation. After the procedure, place the mouse in a supine position with its head raised on the heating pad for recovery. Once the mouse has regained mobility, return it to its original cage. Three days later, the Fast Blue tracer will have reached the ganglia, enabling further experiments such as neural isolation or immunofluorescence. The dye will remain visible for up to 8 weeks.

### RNA-seq library preparation, sequencing, and analysis

After euthanizing the mice, the vagal ganglia were dissected and isolated. RNA from the vagal ganglia was extracted using The RNeasy Mini Kit (Qiagen). The ganglia were placed in a micro glass homogenizer containing Buffer RLT and carefully homogenized to release the RNA. Subsequently, RNA was extracted following the protocol outlined in the RNeasy Mini Kit manual. The purity and concentration of the RNA were measured, with the following standards: RIN > 7.0, 28S/18S > 1.0, m > 20 ng. For samples containing less than 4 μL of RNA, Oligo-dT Primer was added, and the first-strand cDNA was synthesized using the SMART amplification technology. The second-strand cDNA synthesis was performed using KAPA HiFi HotStart DNA Polymerase. The synthesized double-stranded cDNA was purified using Agencourt AMPure XP magnetic beads, and the quality of the amplification product was assessed using an Agilent 2100 chip. Qualified amplification products were subjected to library construction using the Nextera XT DNA Library Preparation Kit. The qualified libraries were transformed into single-stranded circular molecules under certain conditions. The single-stranded circular DNA molecules were replicated by rolling circle replication, forming DNA nanoballs (DNBs) containing over 200 copies of DNA. The obtained DNBs were loaded into the mesh holes on the high-density DNA nanochip using high-density DNA nanoball array technology. Sequencing was performed using the combined probe anchor sequencing (cPAS) technology, with a sequencing read length of PE100. Library construction and sequencing were conducted at BGI (Beijing Genomics Institute). Data analysis was performed using R and BGI platform.

### Statistical analysis

All statistical analyses were performed using R and GraphPad Prism 8.0. Summary data were compared for statistical differences using t-tests or one-way ANOVA. Statistical data are presented as means ± SEM. If no statistical significance was observed, “ns” was used. If there was a statistically significant difference, *p < 0.05 indicates a significant difference; **p < 0.01 indicates a highly significant difference; ***p < 0.001 indicates an extremely significant difference.

---

**Extended Data Fig. 1.**
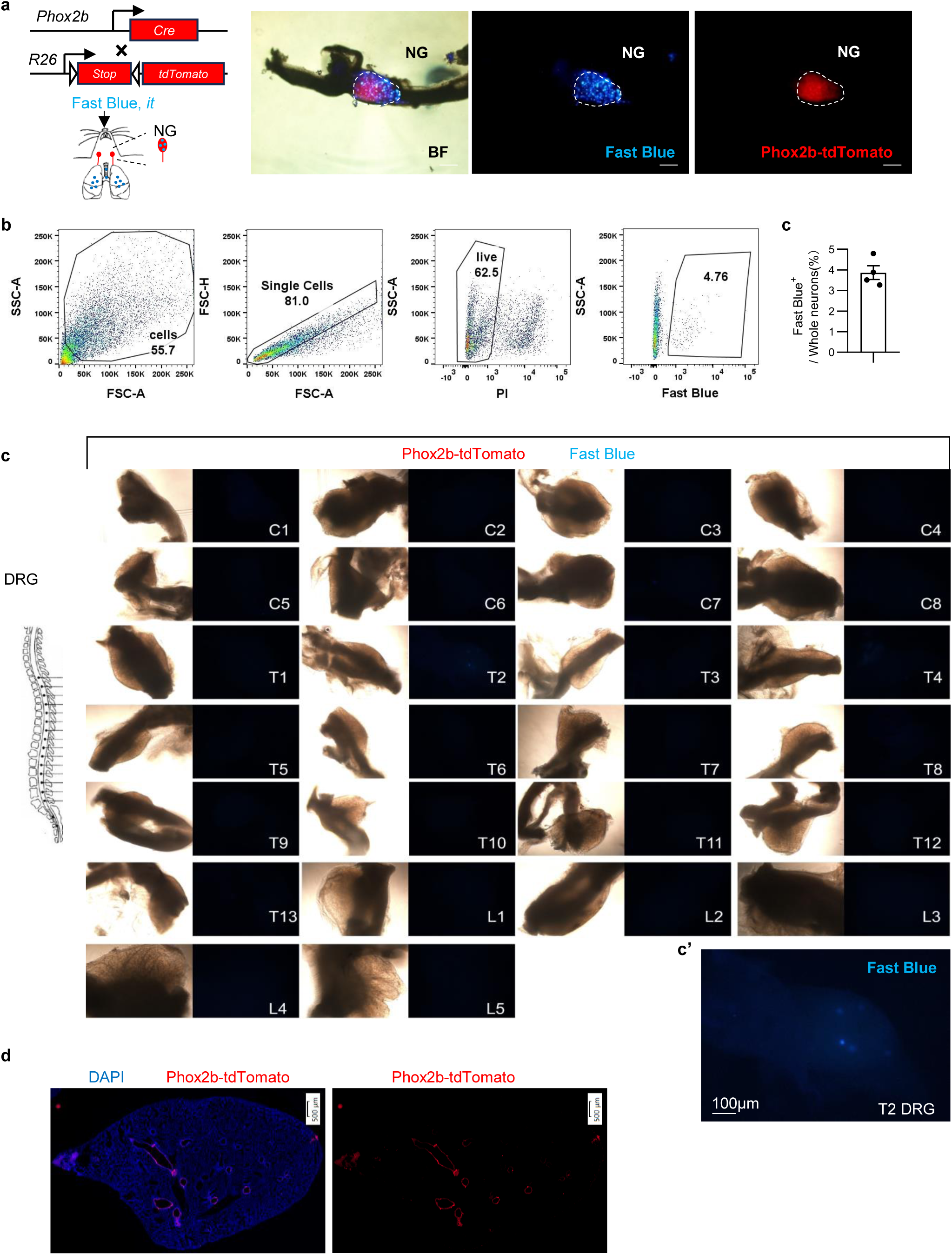
The distribution of sensory neurons projecting to the lung from the NG and DRG. (a) Fast Blue retrograde tracing of sensory neurons projecting to the lung in the nodose/jugular complex of *Phox2b^tdTomato^* mice. (b) Standard flow cytometry gating strategy for Fast Blue retrograde tracing of vagal sensory neurons projecting to the lung (n = 4 mice/group). (c) Fast Blue retrograde tracing of sensory neurons projecting to the lung in the cervical (C1-C8), thoracic (T1-T13), and lumbar (L1-L5) DRGs from adult *Phox2b^tdTomato^* mice. Fast Blue retrograde tracing of sensory neurons projecting to the lung in the T2 DRG of *Phox2b^tdTomato^* mice (C’). (d) Anatomical mapping of vagal sensory neurons within the lung using *Phox2b^tdTomato^* mice. Scale bars: 100 μm in (b) and (c), 500μm in (d).

**Extended Data Fig. 2.**
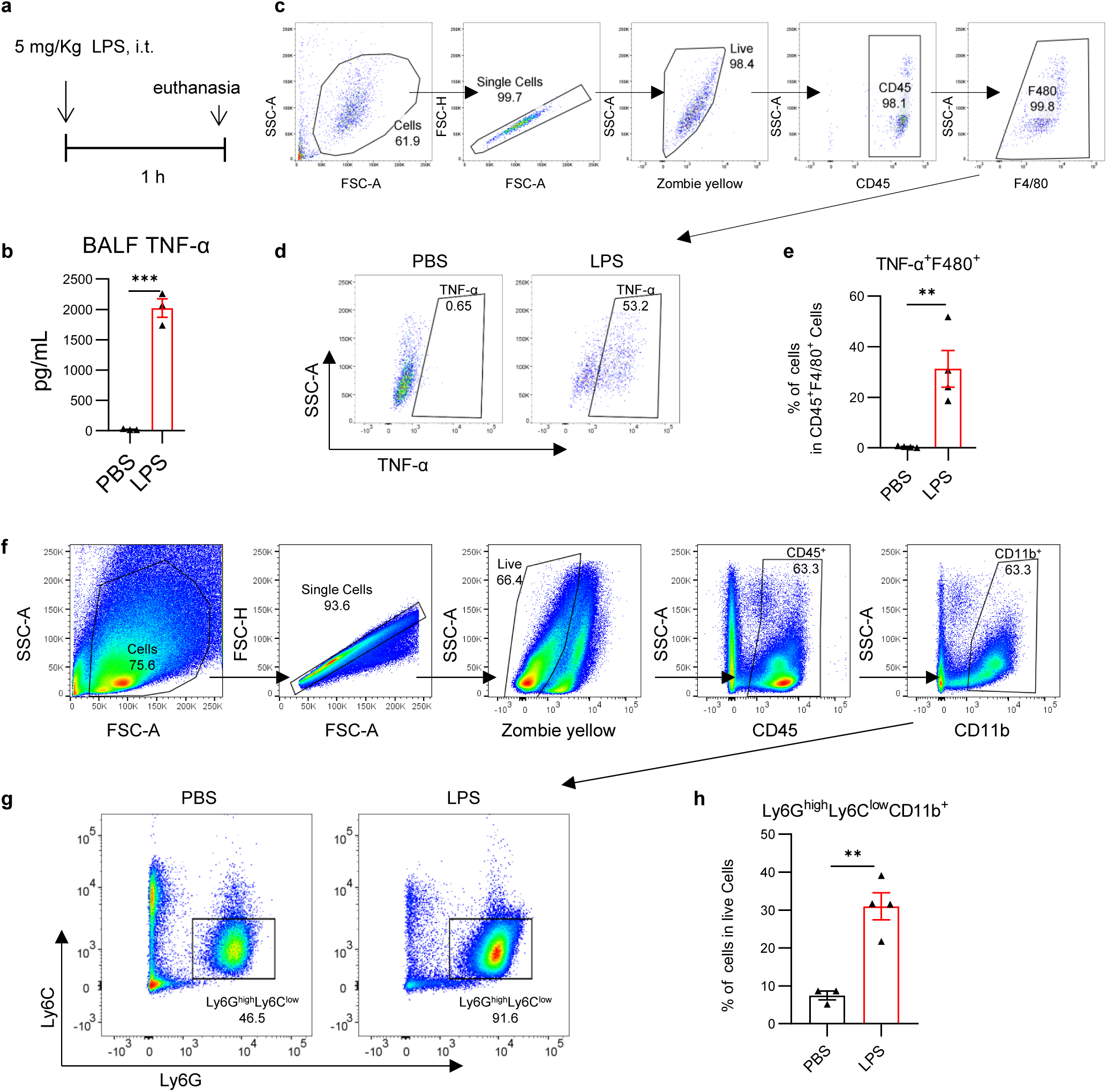
Early pulmonary inflammation induced 1 hour after pulmonary LPS stimulation. (a) Schematic of early pulmonary inflammation induced by pulmonary LPS in mice. (b) TNF-α level in the bronchoalveolar lavage fluid (BALF) of mice 1 hour after pulmonary LPS stimulation (n = 3 mice/group). (c) Standard gating strategies for F4/80^+^ macrophages used in flow cytometry (n = 3 mice/group). (d and e) TNF-α level in F4/80^+^ macrophages in the BALF of mice 1 hour after pulmonary LPS stimulation (n = 3 mice/group). (f) Standard gating strategies for Ly6G^high^Ly6C^low^CD11b^+^ cells (PMNs) used in flow cytometry. The proportion of Ly6G^high^Ly6C^low^ cells among CD11b^+^ cells in lung tissue of mice 1 hour after pulmonary LPS stimulation (n = 3 mice/group). (g and h) The proportion of Ly6G^high^Ly6C^low^CD11b^+^ cells among live cells in lung tissue of mice 1 hour after pulmonary LPS stimulation (n = 3-4 mice/group). Student’s t test in (b), (e) and (h). Mean ± SEM. ns, not significant. *p < 0.05, **p < 0.01, and ***p < 0.001.

**Extended Data Fig. 3.**
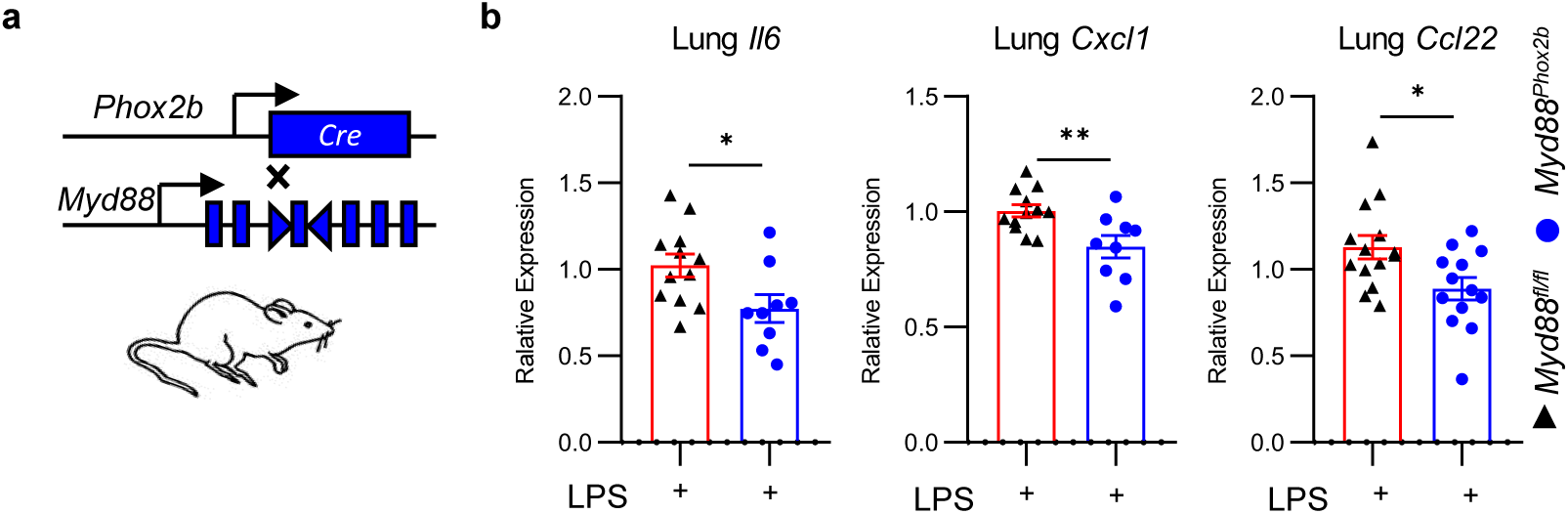
Myd88 deficiency in nodose sensory neurons amplifies LPS-induced pulmonary inflammation. (a and b) The expression levels of *Il6*, *Cxcl1*, and *Ccl22* in lung tissues from *Myd88^Phox2b^* and *Myd88^fl/fl^* mice 1 hour after pulmonary LPS stimulation. Student’s t test in (b). Mean ± SEM. ns, not significant. *p < 0.05, **p < 0.01, and ***p < 0.001.

**Extended Data Fig. 4.**
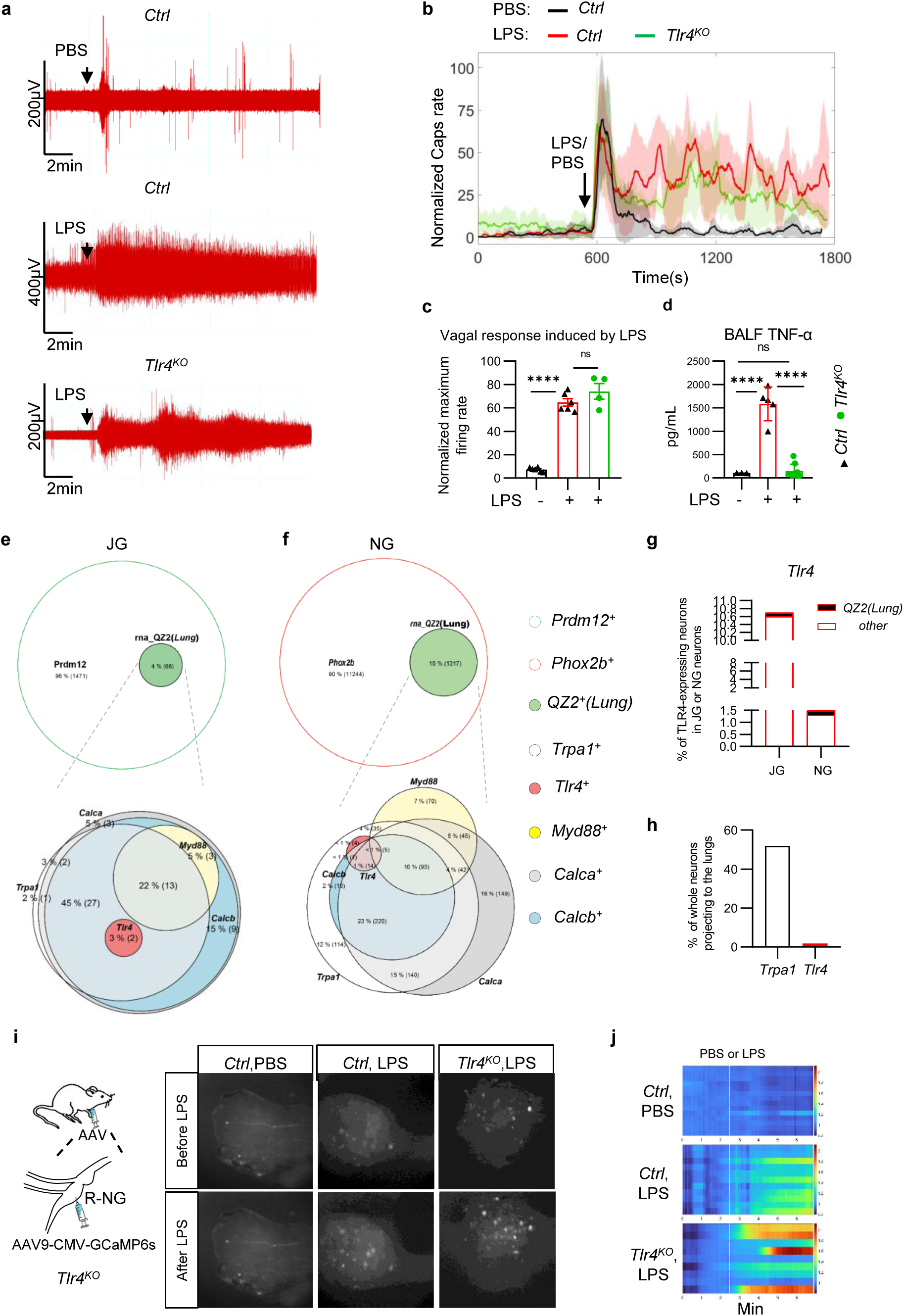
The vagal response induced by pulmonary LPS is independent of TLR4-positive sensory neurons. (a) Vagal responses to pulmonary LPS stimuli within the lung in *Tlr4^KO^*(*Tlr4-/-*) mice and *Ctrl* (*Tlr4+/+*) mice (n = 4-6 mice/group). (b) Statistical analysis of vagal responses to pulmonary LPS stimuli within the lung in *Tlr4^KO^* mice and *Ctrl* mice. *Ctrl*+PBS is represented by gray traces, *Ctrl*+LPS by red traces, and *Tlr4^KO^*+LPS by green traces (n = 4-6mice/group). (c) Quantification of peak responses to LPS in *Tlr4^KO^* mice and *Ctrl* mice (n = 4-6 mice/group). (d) TNF-α levels in the bronchoalveolar lavage fluid (BALF) of *Tlr4^KO^* mice and *Ctrl* mice 1 hour after pulmonary LPS stimulation (n = 4-6 mice/group). (e) The expression of *Trpa1*, *Myd88*, *Tlr4*, *Calca*, and *Calcb* in Prdm12^+^ jugular sensory neurons projecting to the lung. The numbers in the Fig. represent the number of neurons and the percentage of these neurons among all neurons depicted in the subpanel. These data are from a reanalysis of “The full Projection-seq dataset” published by Zhao et al ^32^. (f) The expression of *Trpa1*, *Myd88*, *Tlr4*, *Calca*, and *Calcb* in Phox2b^+^ nodose sensory neurons projecting to the lung. The numbers in the Fig. represent the number of neurons and the percentage of these neurons among all neurons depicted in the subpanel. These data are from a reanalysis of “The full Projection-seq dataset” published by Zhao et al ^32^. (g) The distribution of TLR4-positive neurons in the NG or JG that project to the lungs versus other locations. (h) The ratio of *Trpa1*-positive sensory neurons and *Tlr4*-positive sensory neurons to all sensory neurons projecting to the lung (*QZ2*-positive sensory neurons) ^32^. (i Recording calcium responses in nodose sensory neurons expressing GCaMP6s during intratracheal LPS stimulation in *Tlr4^KO^* mice and *Ctrl* mice. (j) The heat maps depict z-score-normalized fluorescence traces from nodose neurons responding to LPS stimulation in *Tlr4^KO^* mice and *Ctrl* mice. One-way ANOVA in (c) and (d). Mean ± SEM. ns, not significant. *p < 0.05, **p < 0.01, and ***p < 0.001.

**Extended Data Fig. 5.**
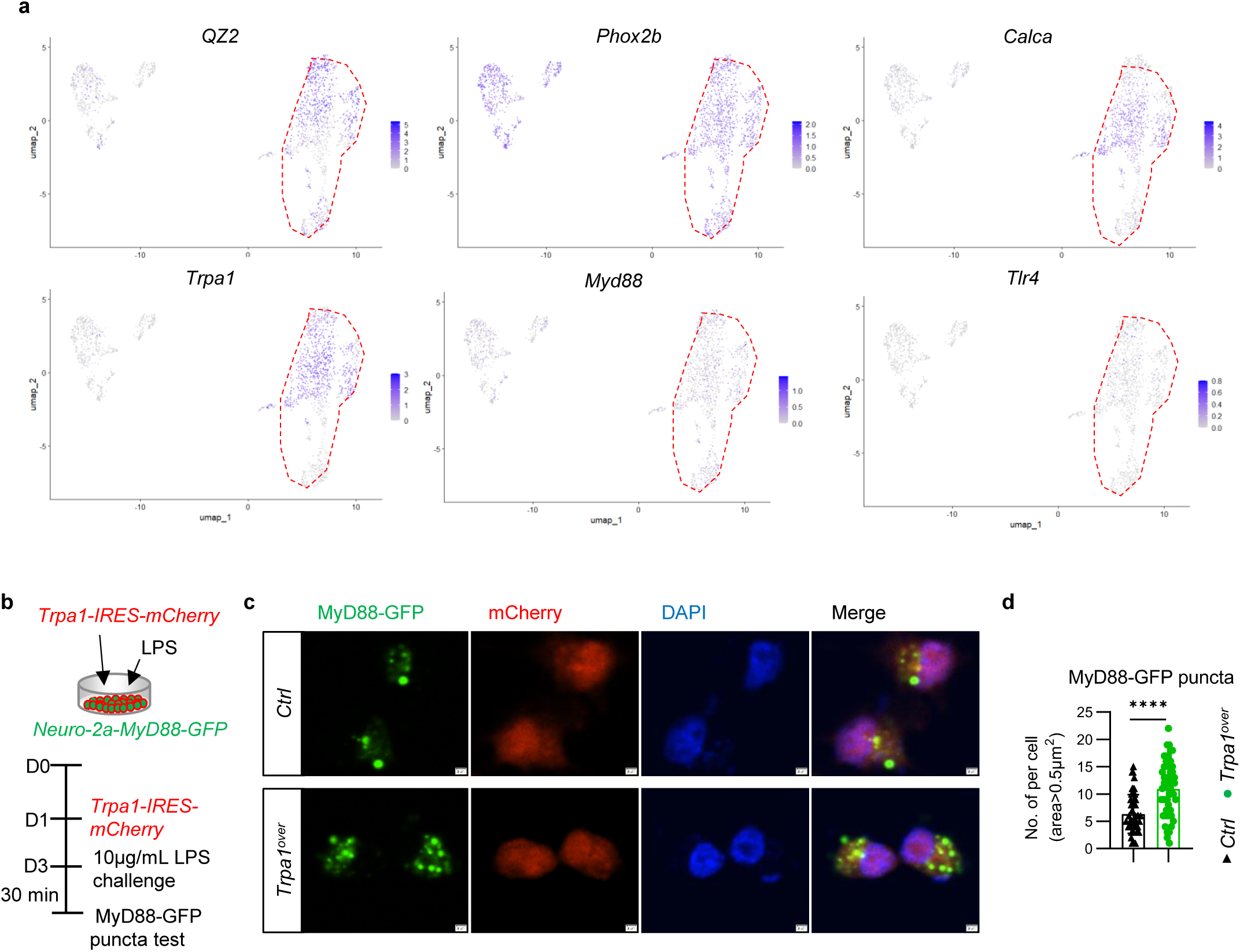
TRPA1 enhances Myd88 oligomerization. (a) The expression of *Phox2b, Calca, Trpa1, Myd88*, and *Tlr4* genes in QZ2^+^ neurons, which are vagal neurons projecting to the lungs. (b) Experimental procedure for detecting Myd88 oligomerization in Neuro-2a cells. (c) MyD88-GFP puncta in Neuro-2a cells overexpressing *Trpa1* (*Trpa1^over^*) and control (*Ctrl*) after 30 minutes of LPS induction. (d) After 30 minutes of LPS induction, the number of MyD88-GFP puncta larger than 0.5 μm^2^ per Neuro-2a cell in Neuro-2a cells overexpressing *Trpa1* (*Trpa1^over^*) and control (*Ctrl*). Student’s t test in (d). Mean ± SEM. ns, not significant. *p < 0.05, **p < 0.01, and ***p < 0.001.

**Extended Data Fig. 6.**
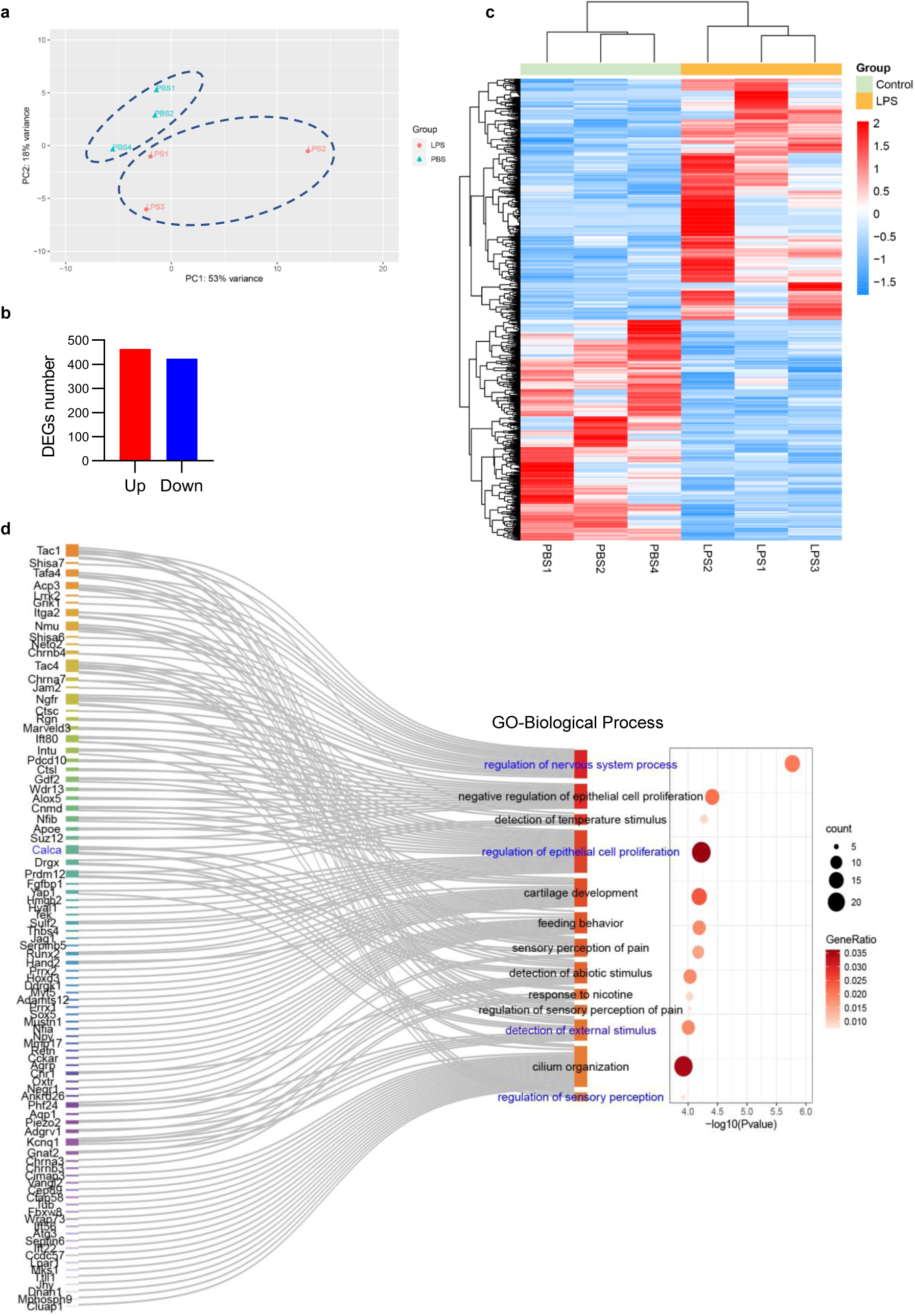
RNA-Seq analysis of vagal sensory neuron response to pulmonary LPS. (a) Principal component analysis (PCA). (b) Quantification of differentially expressed gene numbers in mice treated with PBS vs. LPS. (c) The heat map shows differentially expressed genes identified from RNA-seq data of vagal ganglia in mice treated with PBS vs. LPS. (d) Differential genes can be analyzed by GO biological process clustering, and enriched genes can be visualized using a Sankey diagram.

**Extended Data Fig. 7.**
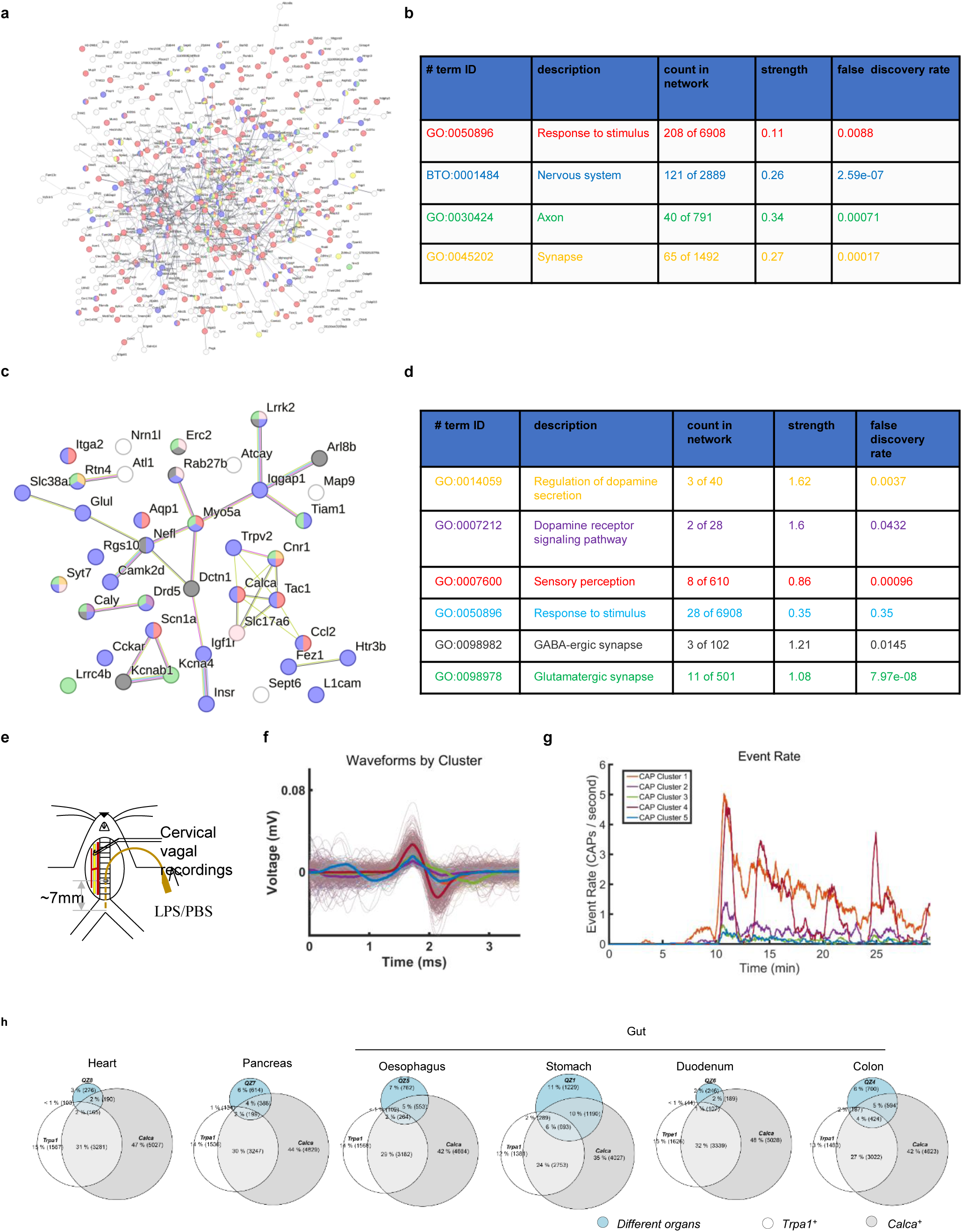
Various types of vagal sensory neuron responses to pulmonary LPS. (a) STRING protein interaction pathway analysis was performed on RNA-seq data from the vagal ganglia of mice treated with PBS vs. LPS. All genes shown are differentially expressed genes (p < 0.05, n = 4 mice/group). (b) Top 4 enriched pathways identified by STRING protein interaction pathway analysis. (c) In anesthetized wild-type mice, lung LPS stimulation was performed via endotracheal intubation, while recording the electrical activity of the cervical vagus nerve. (d) Vagal responses to pulmonary LPS measured by compound action potentials (CAPs). Each colored curve represents the response rate of different CAPs over time. (e) Neural responses of the vagus nerve to LPS as measured by compound action potentials (CAPs). Each colored curve represents the response rate of different CAPs over time. (f and g) STRING protein interaction pathway analysis was performed on synapse-related genes from RNA-seq data. (h) The relationship between vagal sensory neurons projecting to different visceral organs and those expressing *Trpa1*^+^ or *Calca*^+^. The numbers in the Figure represent the number of neurons and the percentage of these neurons among all neurons depicted in the subpanel^32^.

**Extended Data Fig. 8.**
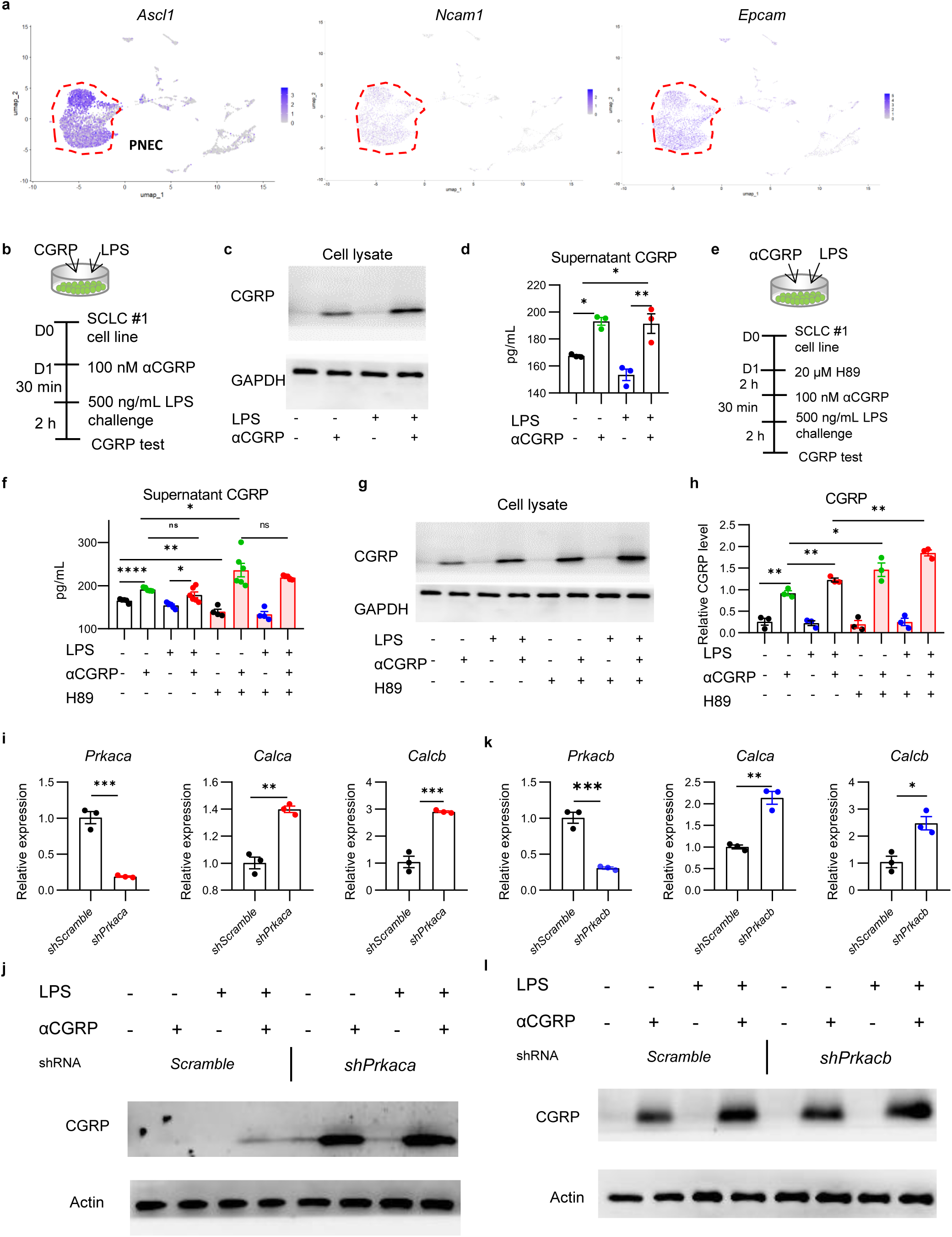
αCGRP can induce excessive CGRP production in PNECs and PNEC responds to αCGRP stimulation via PKA. (a) The expression of *Ascl1, Ncam1*, and *Epcam* in PNECs. These data are from a reanalysis of “PNEC dataset” published by Schappe et al^1^. (b) Experimental timeline for exogenous αCGRP and LPS treatment in SCLC cells. (c and d) The levels of CGRP in both the intracellular compartment (B) and the cell supernatant (C) of SCLC cells stimulated with LPS and αCGRP were measured. (e) SCLC cells were cultured with LPS or αCGRP+LPS with medium or H89. (f) The level of CGRP in the supernatant of SCLC cells treated with LPS or αCGRP+LPS with medium or H89. (g and h) The intracellular level of CGRP in SCLC cells treated with LPS or αCGRP+LPS with medium or H89. (i) The expression of *Prkaca, Calca*, and *Calcb* genes in *Prkaca* knockdown SCLC cells. (j) The intracellular level of CGRP in *Prkaca* knockdown SCLC cells treated with LPS or αCGRP+LPS. (k) The expression of *Prkacb*, *Calca*, and *Calcb* genes in *Prkacb* knockdown SCLC cells. (l) The intracellular level of CGRP in *Prkacb* knockdown SCLC cells treated with LPS or αCGRP+LPS. Student’s t test in (f), (i) and (k). One-way ANOVA in (d) and (h). Mean ± SEM. ns, not significant. *p < 0.05, **p < 0.01, and ***p < 0.001.

**Extended Data Fig. 9.**
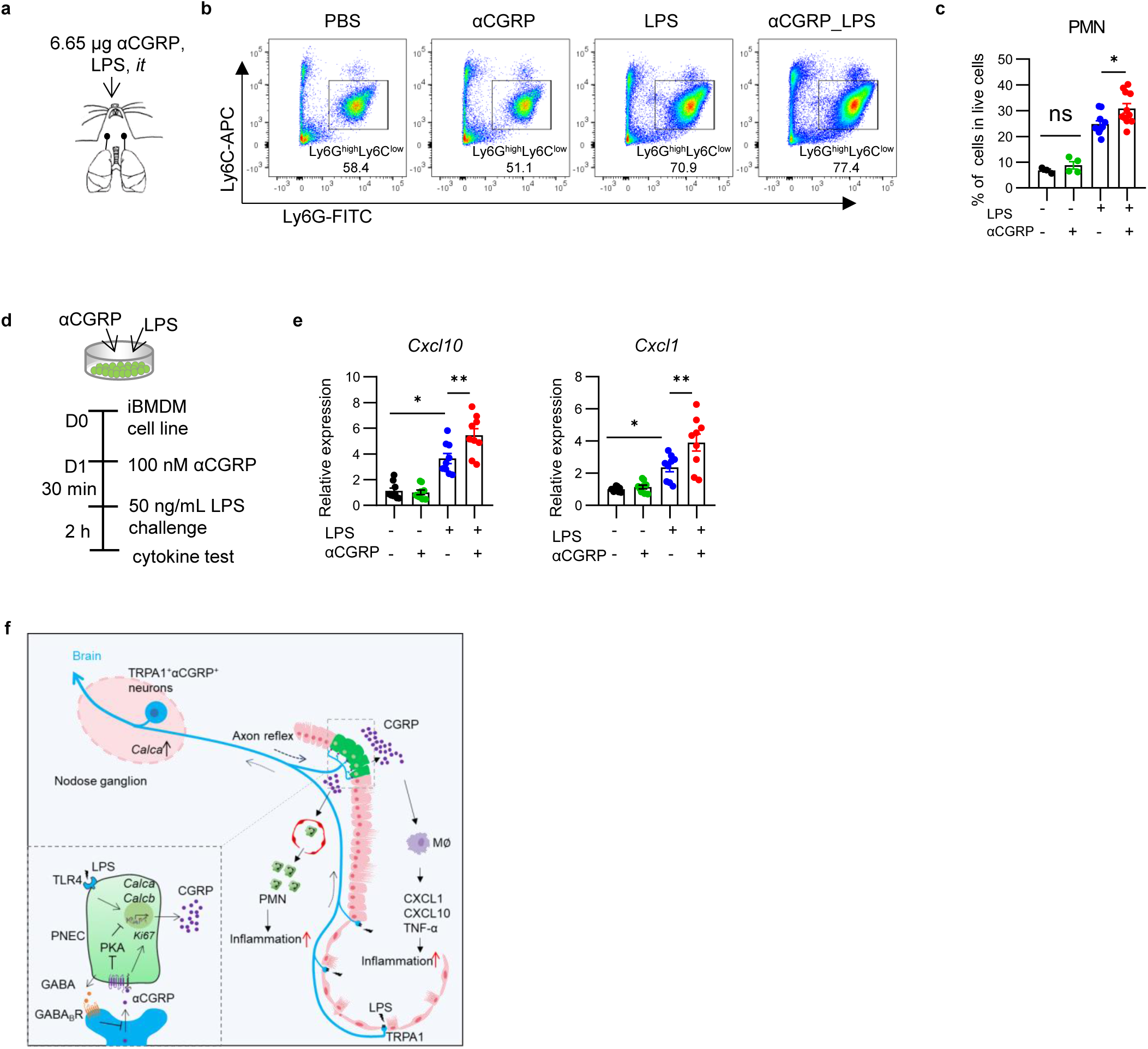
αCGRP promotes the recruitment of immune cells to the lungs. (a) Administration of αCGRP or αCGRP+LPS into the lungs of mice. (b) The proportion of Ly6G^high^Ly6C^low^ cells among CD11B^+^ cells in lung tissue from mice following administration of αCGRP or αCGRP+LPS (n = 4-10 mice/group). (c) The proportion of Ly6G^high^Ly6C^low^CD11B^+^ cells (PMNs) among live cells in lung tissue from mice following administration ofαCGRP or αCGRP+LPS (n = 4-10 mice/group). (d) Administration of αCGRP or αCGRP+LPS in iBMDM cells. (e) Expression levels of *Cxcl1* and *Cxcl1*0 in iBMDM cells cultured with αCGRP, LPS, or αCGRP+LPS. (f) Graphical abstracy of αCGRP^+^TRPA1^+^ nodose sensory neurons regulating PNECs via αCGRP. One-way ANOVA in (c), (e) and (h). Mean ± SEM. ns, not significant. *p < 0.05, **p < 0.01, and ***p < 0.001.

**Extended Data Fig. 10.**
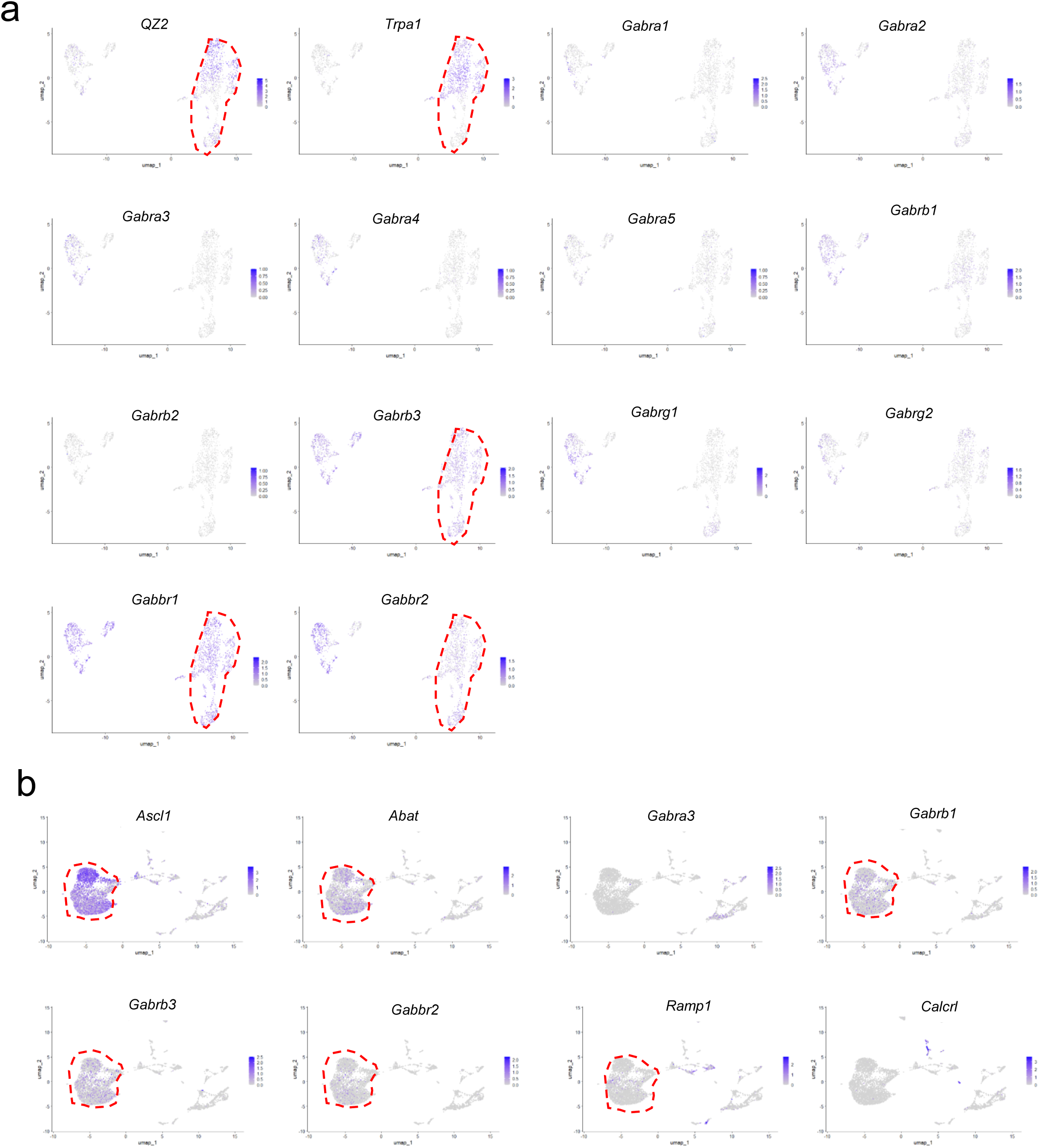
GABA receptors in TRPA1^+^ lung-innervating neurons and GABA/CGRP receptors in PNECs. (a) The expression of GABA receptor genes in QZ2^+^ neurons, which are vagal neurons projecting to the lungs. These data are from a reanalysis of “The full Projection-seq dataset” published by Zhao et al (Zhao et al., 2022). (b) The expression of GABA and CGRP receptors in PNECs. These data are from a reanalysis of “PNEC dataset” published by Schappe et al (Schappe et al., 2024).

## Notes

### Competing Interest Statement

The authors have declared no competing interest.

